# The Medicines for Malaria Venture Malaria Box contains inhibitors of protein secretion in *Plasmodium falciparum* blood stage parasites

**DOI:** 10.1101/2022.05.01.490233

**Authors:** Oliver Looker, Madeline G. Dans, Hayley E. Bullen, Brad E. Sleebs, Brendan S. Crabb, Paul R. Gilson

## Abstract

*Plasmodium falciparum* parasites which cause malaria, traffic hundreds of proteins into the red blood cells (RBCs) they infect. These exported proteins remodel their RBCs enabling host immune evasion through processes such as cytoadherence that greatly assist parasite survival. As resistance to all current anti-malarial compounds is rising new compounds need to be identified and those that could inhibit parasite protein secretion and export would both rapidly reduce parasite virulence and ultimately lead to parasite death. To identify compounds that inhibit protein export we used transgenic parasites expressing an exported nanoluciferase reporter to screen the Medicines for Malaria Venture Malaria box of 400 anti-malarial compounds with mostly unknown targets. The most potent inhibitor identified in this screen was MMV396797 whose application led to export inhibition of both the reporter and endogenous exported proteins. MMV396797 mediated blockage of protein export and slowed the rigidification and cytoadherence of infected RBCs - modifications which are both mediated by parasite-derived exported proteins. Overall, we have identified a new protein export inhibitor in *P. falciparum* whose target though unknown, could be developed into a future anti-malarial that rapidly inhibits parasite virulence before eliminating parasites from the host.

**Synopsis:** *Plasmodium falciparum* exports proteins into its host cell to perform a myriad of functions required for survival. We adapted an assay to screen for small molecules that inhibit protein secretion and export. Screening the 400-compound Medicines for Malaria Venture (MMV) Malaria Box uncovered several potential export inhibitors. The most promising of these compounds, MMV396797, blocked protein export at the parasite and reduced host rigidification and cytoadherence, two functions which are mediated by exported proteins.

## Introduction

Malaria caused by infection with *Plasmodium* parasites remains a major global health burden and triggered over 241 million cases in 2020, tragically resulting in an estimated 627,000 deaths ^1^. This 12% recent increase in malaria associated deaths has been attributed to service disruptions due to the COVID-19 pandemic ^1^. *Plasmodium falciparum* is the cause of most of these deaths and is predominantly found throughout sub-Saharan Africa. Antimalarial drug treatment remains the only means of clearing parasite infection and artemisinin-based therapies are the gold standard. Alarmingly however, mutations in *P. falciparum* parasites that confer resistance to these current frontline therapies are being detected in many regions of the world after having first appeared in the Greater Mekong Subregion of Southeast Asia ^2–5^. To overcome resistance to artemisinin monotherapy, these fast acting but rapidly eliminated compounds have been partnered with slower acting but more persistent compounds including lumefantrine, mefloquine, piperaquine, amodiaquine, sulfadoxine and pyrimethamine ^6^. Parasite resistance to the partner compounds is now becoming increasingly common ^1^, highlighting the importance of novel antimalarial discovery.

To develop new antimalarial medicines, millions of compounds have been screened for anti-parasite activity and tens of thousands of potent, parasite killing compounds have been identified ^7–9^. To accelerate the identification of drug targets and to engender great interest and collaboration in translational neglected disease research, the Medicines for Malaria Venture (MMV) have made freely available several compound libraries containing antimalarial and antipathogen compounds ^10–12^. As compounds in drug libraries were originally screened against asexual blood stages for inhibition of growth, additional phenotypic screens of these libraries have been conducted to identify compounds that inhibit other stages of the lifecycle or target specific enzymes or biological processes ^13–23^. Example phenotypic screens include the recent examination of MMV’s Malaria and Pathogen boxes for inhibitors of parasite egress and invasion of RBCs ^24,25^. It was necessary to independently validate inhibition data from these screens using additional cell-based approaches such as live cell imaging to ensure the compounds acted specifically against egress and invasion as many of the compounds appeared to act against housekeeping functions generally required during the rest of the cell cycle ^24^.

One source of druggable targets is the parasite’s protein trafficking system and inhibitors of PfPI4KIII*β* which block Golgi apparatus to plasma vesicular membrane trafficking have been identified ^26–29^. As typical eukaryotes, *Plasmodium* parasites traffic proteins via the endoplasmic reticulum (ER) *en route* to specialised parasite compartments including the algal-derived apicoplast, the apical secretory organelles (the rhoptries, micronemes and dense granules), and the specialised membranous sac (parasitophorous vacuole membrane (PVM)) encasing the parasite within the RBC ^30^. Parasites also export proteins beyond this encasing membrane and into the RBC cytosol, whereupon they serve many highly specialised roles to support parasite growth and immune evasion. This is a very significant function of the parasite trafficking network, as an estimated 400-500 proteins, representing nearly 10% of their proteomes, are exported into their host RBC ^31–33^. Most of these exported proteins contain a PEXEL motif which serves as a proteolytic cleavage site for the ER-resident protease plasmepsin V ^34,35^. PEXEL cleavage licences proteins for export via the Plasmodium Translocon for EXported proteins (PTEX) which resides in the PVM and serves to unfold and extrude PEXEL and PEXEL-negative exported proteins (PNEPs) into the RBC compartment ^36–39^. Inhibitors of plasmepsin V have been developed that prevent cleavage and therefore export of PEXEL proteins and result in parasite death ^40–42^. Theoretically the most druggable component in PTEX is the unfoldase AAA+ ATPase chaperone HSP101. PTEX’s other core subunits PTEX150 and EXP2, are the structural and membrane spanning components of the protein translocon, respectively ^36,37,43,44^. The importance of protein export for parasite survival is demonstrated by knocking-down expression of PTEX proteins which results in rapid parasite death ^37,39^. HSP101 appears to first recognise protein cargos in the ER and accompany them to PTEX for translocation into the RBC indicating HSP101 inhibitors might result in protein cargoes becoming trapped in the ER and parasitophorous vacuole (PV) ^45,46^.

Although export of parasite proteins into the RBC plays a major role in the virulence of *P. falciparum,* much remains to be resolved about the protein export pathway. The discovery of small molecule inhibitors of protein export would aid in unravelling this complex and essential process. Specifically, many exported proteins are devoted to modifying the infected RBC (iRBC) surface. These modifications include: the insertion of membrane channels for nutrient uptake ^47–50^; changes to iRBC rigidity ^51^ and display of specialised surface molecules (*Pf*EMP1s) which enable the iRBC to cytoadhere to the vascular endothelium. Cytoadherance thereby removes iRBCs from circulation and prevents their passage through the spleen where they would be removed by the immune system ^52^. Exported proteins exposed on the iRBC surface can also recruit neighbouring uninfected RBC to form ‘rosettes’ which along with cytoadhering iRBCs, can occlude blood flow in capillary beds leading to organ damage ^53^. This can result in both cerebral and placental malaria which are extremely dangerous malaria sequelae that can cause death and stunting, respectively ^54,55^.

Identifying small molecules that inhibit protein export would therefore have the dual effect of not only causing death of the parasite, but also rapidly reducing the harm parasites could do to the host. Although PfPI4KIII*β* inhibitors have undergone pre-clinical development and recently phase 1 clinical trials ^29,56^, and greatly improved plasmepsin V inhibitors have recently been evaluated ^57^, we still need to discover compounds with new targets and mechanisms of action. We therefore sought to screen the MMV Malaria Box for inhibitors of parasite protein secretion into the PV (ie. compounds that block steps in the export pathway inside the parasite), and export into the host RBC (ie. compounds that block export through PTEX) by monitoring changes in the location of an exported bioluminescent reporter protein within the iRBC. Our screen identified 14 inhibitors from the 400-compound library that significantly blocked export (by >25%) and subsequently caused parasite death, however a counter screen indicated 10 of these compounds inhibited the activity of the bioluminescent reporter and/or were PfATP4 inhibitors and did not inhibit protein export directly. Hits that passed the counter screen were further analysed by microscopy and it was determined that MMV396797 was a robust export inhibitor of both the bioluminescent reporter and endogenous proteins. Importantly, treatment with MMV396797 also reduced virulence related rigidification and cytoadherence of iRBCs demonstrating that inhibiting protein export with small molecule inhibitors not only results in parasite death, but also rapidly reduces the ability of the parasite to cause harm to the host. Collectively these data strongly support the notion of identifying and developing small molecule inhibitors of protein export as novel antimalarials with the dual purpose of killing the parasite and reducing parasite virulence.

## Results

### Establishing and Validating the Assay

To identify small molecules that inhibit protein export, we first sought to adapt a previously reported assay to quantify protein secretion and export in a higher throughput format ^58^. The assay uses *Plasmodium falciparum* trophozoite stage parasites transfected with a reporter protein cargo comprising the N-terminal 113 amino acids of the exported PEXEL protein *Pf*Hyp1 [PF3D7_0113300], fused to nanoluciferase (Hyp1-Nluc)(Figure 1A,B)^58^. Hyp1-Nluc is first imported into the ER inside the parasite compartment and then secreted into the PV surrounding the parasite before being exported into the RBC. The simplified workflow of the export assay was to treat Hyp1-Nluc trophozoite (20-24 hpi, hours post invasion) infected RBCs (iRBCs) with 10 µM of the MMV Malaria Box compounds diluted in DMSO for 5 hours. Treated parasites were subsequently split into three aliquots. The first aliquot was treated with equinatoxin (Eqt) to permeabilise the RBC membrane and release the contents of the RBC cytoplasm (Figure 1A). The second aliquot was permeabilised with equinatoxin and saponin (Sap) to release the contents of the RBC and the PV (Figure 1A). The final aliquot was treated with the non-ionic detergent Igepal CA-630 (Det) in hypotonic buffer to permeabilise all the membranes (Figure 1A). All lysis buffers contained the membrane-impermeable nanoluciferase substrate NanoGlo, which reacts with the Hyp1-Nluc reporter released during differential lysis to produce bioluminescence signals proportional to the amount of Hyp1-Nluc present in each compartment (Figure 1A). The three lysates were normalised against parasites treated with a non-lysing phosphate buffer to account for background lysis. To account for minor differential activity of nanoluciferase in each of the lysis buffers, the activity of equal amounts of recombinant nanoluciferase was measured in each of the buffers to normalise Nluc activity to each of the parasite compartments. Subtraction of the saponin bioluminescence signal (yellow bar) from the detergent bioluminescence signal (blue bar), revealed the amount of Hyp1-Nluc trapped in the parasite compartment. The anticipated levels of Hyp1-Nluc in each compartment are graphically represented at the bottom of Figure 1A.

**Figure 1.**
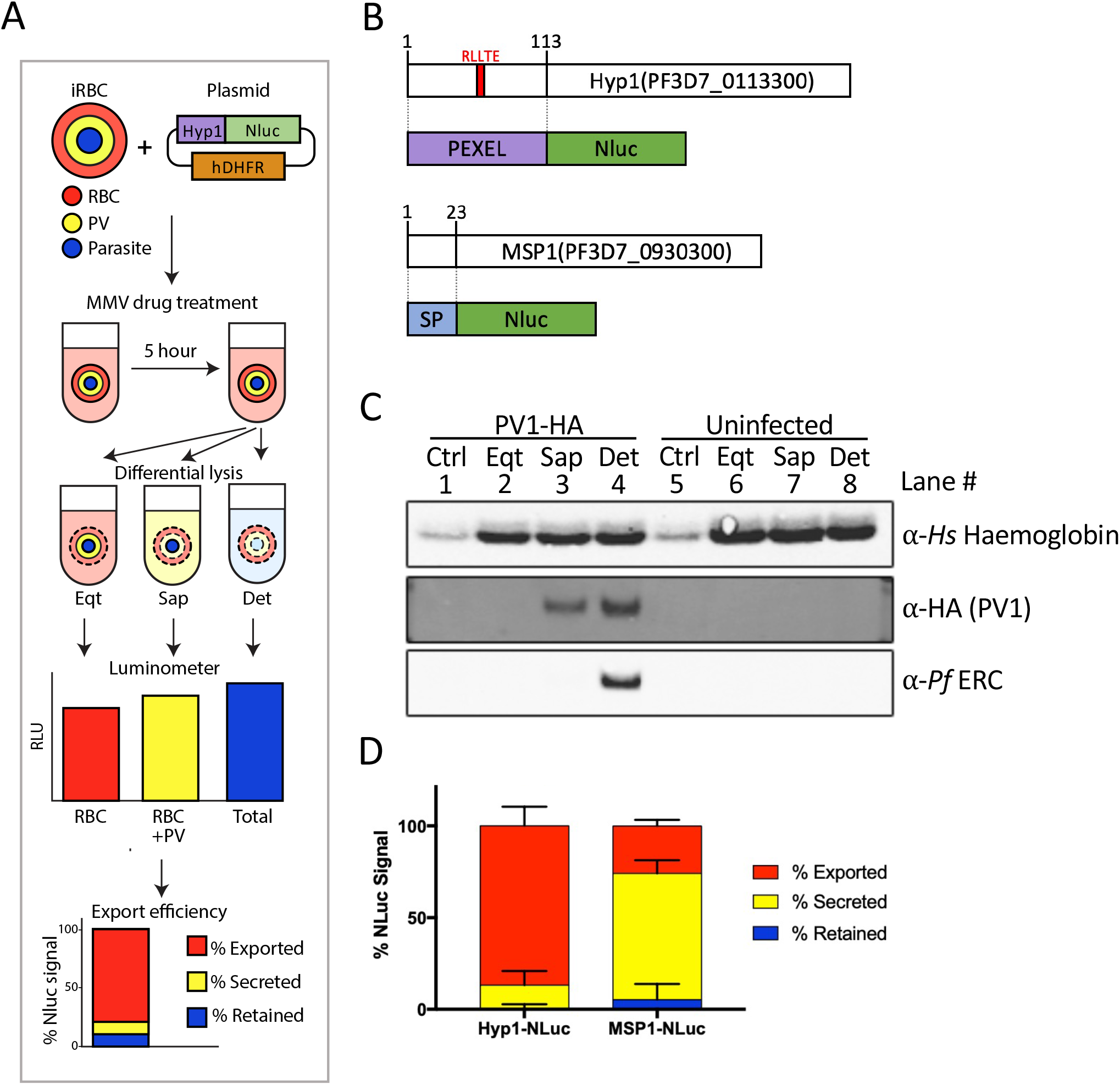
A nanoluciferase based assay for screening inhibitors of protein export. **A)** A simplified schematic diagram summarising the adapted export assay. Whole infected RBCs were differentially lysed using 3 separate buffers containing either Equinatoxin (Eqt), saponin (Sap) and Equinatoxin or Igepal CA-630 (Det) in a hypotonic buffer to lyse the RBC, PV and parasite membranes, respectively. Bioluminescence signals from nanoluciferase released during lysis of each compartment was measured where signal from the RBC, PV and parasite are displayed in red, yellow and blue, respectively. **B)** Diagrammatic representation of the exported Hyp1-Nluc and secreted MSP1-Nluc constructs. The Hyp1-Nluc construct contains a PEXEL sequence (red) within the first 113 amino-acids of Hyp1 (purple), fused to nanoluciferase. The MSP1-Nluc construct contains the signal peptide (SP, blue) from MSP1 fused to nanoluciferase. **C)** Western blot analysis of the export assay lysis conditions. PV1-HA-glmS infected and uninfected RBCs were treated with buffers from the export assay described in A). Blots were labelled with anti-*Hs* hemoglobin, anti-HA (to detect PV1-HA) and anti-*Pf*ERC primary antibodies to label proteins from the RBC, PV and parasite fractions, respectively. Buffers used are labelled as follows: Crtl = Control Tris-phosphate buffer; Eqt = Equinatoxin buffer; Sap = Saponin buffer; Det = Hypotonic Igepal CA-630 buffer. **D)** Export assay analysis of Hyp1-Nluc or MSP1-Nluc fusion protein location. 20-24 hpi Hyp1-Nluc parasites were selectively lysed as described in Figure 1A). Nanoluciferase activity in each compartment was calculated from 2 technical replicates from each of 3 biological repeats.

Prior to performing the screen, we first validated the permeabilisation conditions of the assay by monitoring release of haemoglobin (as a measure of RBC cytoplasm release), PV-resident protein PV1 (a soluble marker of PV release, detectable in a PV1-HA parasite line ^59^), and *Pf* ERC (as a measure of release from the parasite ER)^60^. As expected, western blots of proteins solubilised by these treatments indicated that equinatoxin only releases haemoglobin, demonstrating that only the RBC membrane is lysed under these conditions (Figure 1C, lane 2 (Eqt)). Furthermore, saponin treatment releases the contents of the PV as revealed by the presence of PV1 (Figure 1C, lane 3 (Sap)), and inclusion of Igepal CA-630 in hypotonic buffer released the contents of the parasite’s internal organelles as demonstrated by the presence of *Pf* ERC (Figure 1C, lane 4 (Det)). We then performed the export assay with the exported Hyp1-Nluc reporter, as well as an MSP1-Nluc reporter (containing a classic ER-signal sequence that is secreted into the PV ^58^) and quantified the nanoluciferase signal in each compartment. There is clear export of the Hyp1-Nluc reporter to the erythrocyte cytoplasm (Figure 1D, red bar), compared with MSP1-Nluc which primarily localises to the PV space (yellow bar). We note that although the nanoluciferase signal for MSP1-Nluc was primarily detected in the PV, we also observed 25.9 ± 3.3 % nanoluciferase signal in the exported RBC fraction. Our western blot data (Figure 1C) show no PV1 signal in the RBC fraction and it is therefore likely that some unwanted reporter-specific export of MSP1-Nluc occurs, consistent with a previous study using this cell line ^58^.

### Screening the MMV Malaria Box for inhibitors of Export

After confirming that the export assay buffers worked as expected, we validated the assay using known secretion inhibitors. Specifically, we used Brefeldin A which blocks vesicular trafficking between the ER and Golgi ^61^, and Torin 2. The latter is a known trafficking inhibitor which likely targets *Plasmodium* phosphatidylinositol 4-kinase (PI4KIII*β*), blocks vesicular trafficking at the Golgi and has also been found to deplete proteins at the PVM ^27,28^. These control compounds caused accumulation of Hyp1-Nluc in the parasite (Brefeldin A) and in the parasite and PV (Torin 2) (Figure 2A). We also used commercial antimalarials artemisinin and chloroquine as negative control compounds that have mechanisms of action independent of protein export. Even though all compounds appeared to reduce parasite growth, as evident by a strong reduction in the total bioluminescence of the iRBCs compared to the DMSO vehicle, the export blocking effects of brefeldin A and Torin 2 appeared to be specific because these effects were not observed with the antimalarial negative control compounds (Figure 2A).

**Figure 2.**
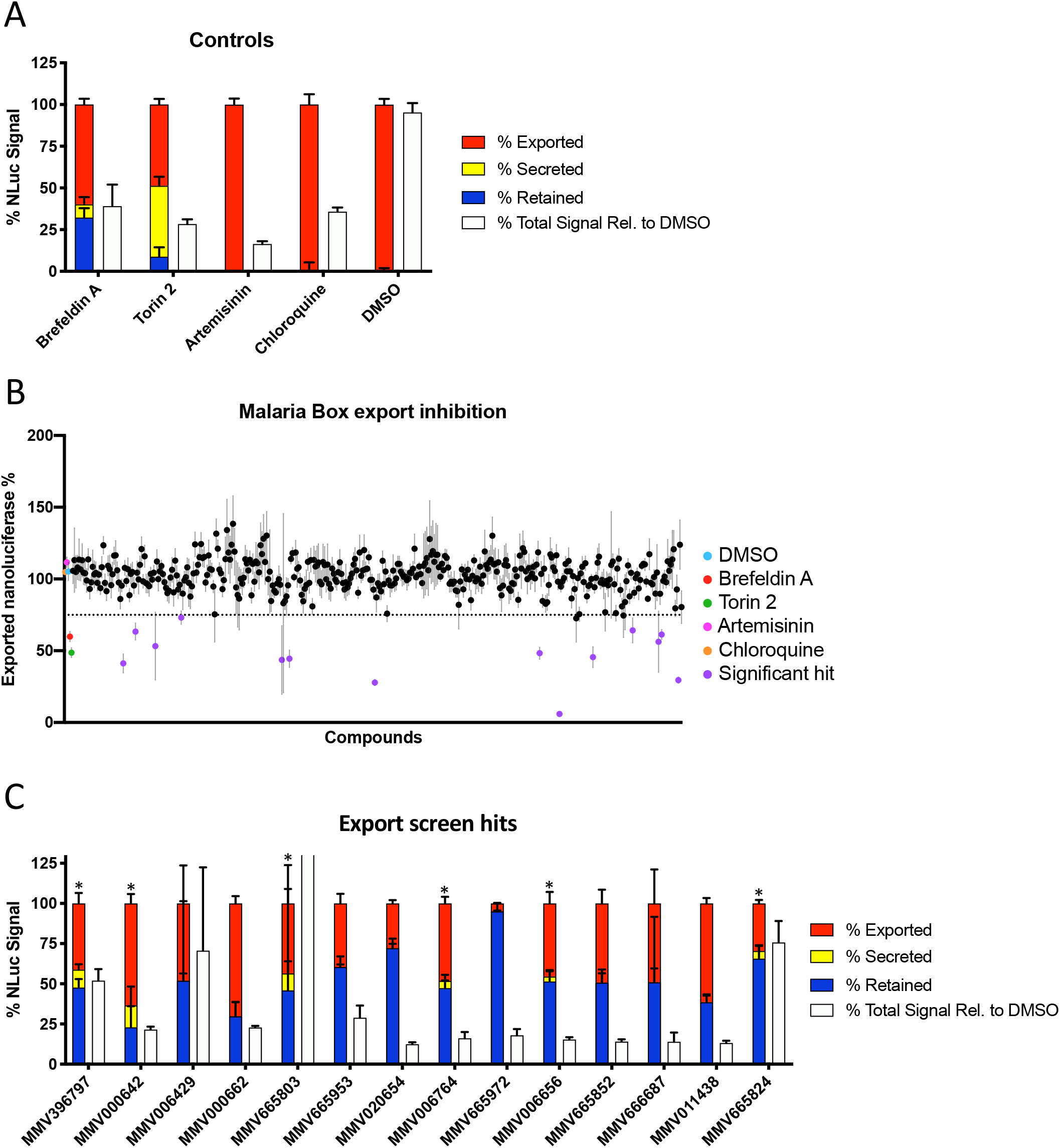
Screening the MMV Malaria Box reveals inhibitors of protein export. **A-C)** Screening of export controls by export assay. Hyp1-Nluc parasites at 20-24 hpi were treated for 5 hours with A,B) Brefeldin A (10 μM), Torin 2 (10 μM), Artemisinin (10 μM), Chloroquine (10 μM), DMSO (0.1 %) or B,C) MMV Malaria Box compounds (10 μM). The export assay was performed as described in Figure 1A. Total cellular nanoluciferase signal normalised to the DMSO control group is shown in white in A,C). Data was collected over multiple replicates for each control treatment as follows: DMSO (n = 44), Artemisinin (n = 20), Chloroquine (n = 20), Brefeldin A (n = 16) and Torin 2 (n = 20) and MMV compounds (n = 4). Compounds were considered hits if they significantly inhibited export when compared to the DMSO control (one-way ANOVA, p < 0.05). Data are presented as mean percentage ± SEM. The dotted line in panel B) represents 75% exported signal. Asterisks in panel C) mark compounds with signal in the PV.

The validity of high throughput screens is often tested by calculating a Z-score (Table S1), showing how many standard deviations each compound-treated population is from a baseline population ^62^. Due to the time-sensitive nature of protein export, standard deviations between biological replicates with the Hyp1-Nluc parasites were relatively large. Paired with the non-binary readout of signal from each of 3 compartments, Z-scores were deemed inappropriate for this assay. We instead decided to proceed with the screen of the 400-compound MMV Malaria Box with rigorous follow up of hits through subsequent microscopy and phenotypic tests as further validation of the export screen. Using the assay, we screened the entire 400-compound MMV Malaria Box, identifying fourteen compounds that statistically significantly inhibited export into the RBC compartment compared to a DMSO vehicle control (one-way ANOVA, p < 0.05) (Figure 2B and Table S2). A further breakdown of the hit compounds into the relative amounts of the reporter in each compartment indicated an increased proportion of Hyp1-Nluc signal retained in the parasite compared to the exported fraction, while the PV signal only slightly increased for six compounds (Figure 2C, asterisks).

### Counter screening the Malaria box reveals inhibitors of nanoluciferase and of protein export

To account for potential false positive results, we performed a counter screen of the whole library to determine if any compounds inhibited nanoluciferase activity. As there was insufficient material remaining in our original Malaria Box to perform the counter screen, we used a newer version of the Malaria Box. This differed slightly from the original box in that it did not contain the hit compound MMV020654 and we were therefore only able to retest 13 of our original 14 export inhibitors. To perform the counter screen, we analysed the extent of nanoluciferase activity inhibition in Hyp1-Nluc parasites with or without Igepal CA-630 as the bioluminescence output of luciferase-based reactions can be affected by the presence of non-ionic detergents ^63^ (Figure 3A, Table S3). Of the 13 primary hit compounds, 10 inhibited nanoluciferase itself to some degree, with 6 compounds inhibiting nanoluciferase by more than 50% at 10 µM (Figure 3A). Additionally, 6 of these compounds inhibited nanoluciferase to a lesser extent in the presence of detergent, which would have resulted in artefactual inhibition of export using our assay (Figure 3A, unpaired t-test, p < 0.05). Taking the entire Malaria Box into account, 14 of 400 compounds inhibited nanoluciferase greater than 50% at 10 µM (Figure S1A), and of these, 11 were statistically significantly less inhibitory in the presence of Igepal CA-630 (Figure S1B, unpaired t-test, p < 0.05). There are however several compounds that were equally inhibitory with and without detergent, indicating multiple mechanisms of nanoluciferase inhibition amongst the set of compounds. Notably MMV396797, MMV665953 and MMV666687 did not inhibit nanoluciferase activity with or without Igepal CA-630 (underlined, Figure 2A), indicating they are likely blocking protein export and not nanoluciferase. The chemical structures of these compounds are shown in Figure 3B. Each of the identified compounds belong to structurally distinct chemical classes. MMV396797 is a pyralopyrimidine and this chemotype shares distinct similarities to kinase-like inhibitors. MMV665953 belongs to the diaryl urea structural class and shares close similarity to the diaryl urea, MMV665852. MMV666687 is from the O-acyl oxime structural class and it is possible the O-acyl oxime is reduced or hydrolysed by metabolic processes in the parasite to give an amidine or amidine oxime by product. It is unknown whether either amidine metabolite is the active constituent of MMV666687, but a variety of amidine containing compounds have been shown to have a wide range of antiparasitic activities ^64^.

**Figure 3.**
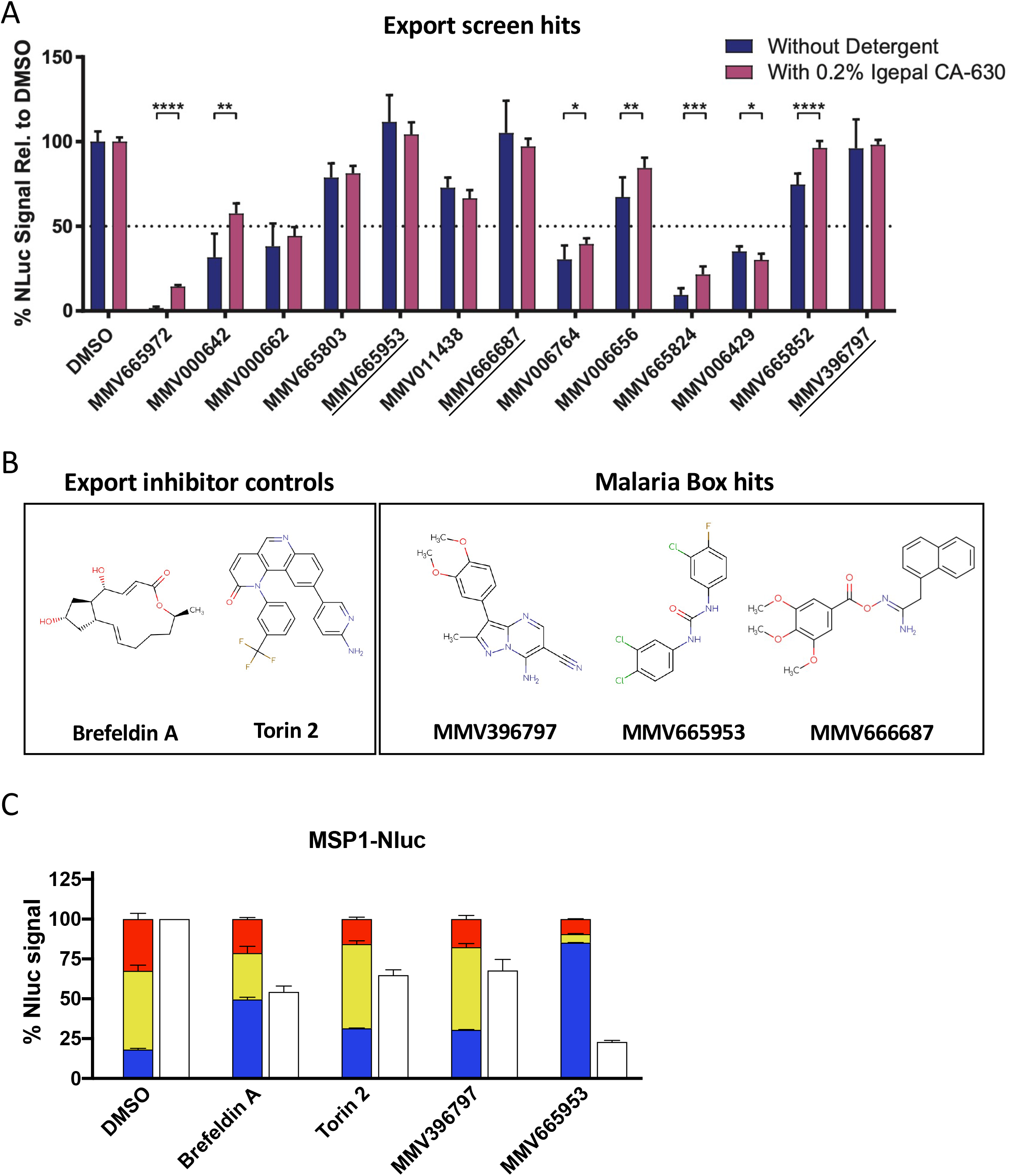
Counter screening the MMV malaria box reveals inhibitors of nanoluciferase activity. **A)** Counter screen of the significant hits from the Malaria Box in the presence or absence or Igepal CA-630. Parasite lysate from Hyp1-Nluc parasites was treated with MMV Malaria Box compounds (10 μM) or DMSO (0.1 %) and total nanoluciferase signal was measured in the presence or absence of Ipegal CA-630. Mean signal ± SEM is presented for the lead hits from Figure 2B,C. The statistical difference between the mean for each treatment with or without Igepal CA-630 was calculated using an unpaired t-test (* p < 0.05, ** p ≤ 0.01, *** p ≤ 0.001, **** p ≤ 0.0001). 50% signal inhibition is marked with a dotted line. **B)** Chemical structures of the known secretion inhibitors Brefeldin A and Torin 2, as well as the Malaria box hits MMV396797, MMV665953 and MMV666687. Structures are coloured as follows: oxygen (red); nitrogen (blue); chlorine (green); fluorine (brown). **C)** Inhibition of MSP1-Nluc secretion. Parasites expressing an MSP1-Nluc fusion protein were synchronised to 20-24 hpi and treated for 5 hours with DMSO, Brefeldin A, Torin 2, MMV396797 or MMV665953. Cells were selectively lysed with equinatoxin II, saponin or Igepal-630 to release protein exported into the RBC (red), secreted into the PV (yellow) and retained within the parasite (blue). Total cellular nanoluciferase activity is presented in white. All values are presented as mean percentage ± SEM. Experiments in A) and C) were performed in technical duplicates over 3 biological replicates.

### MMV396797 and MMV665953 block secretion of parasitophorous vacuole proteins

The initial export screen that identified MMV396797, MMV665953 and MMV666687 was completed on the Hyp1-Nluc parasite line, which specifically investigates the effect of compounds on protein export into the RBC via PTEX at the PVM. However, given that Hyp1-Nluc is trafficked through the ER the prior to presentation at the PVM, blocking the export of this protein across the PVM also effects protein trafficking within the parasite as can be seen by the large blue bars (Nluc trapped in the parasite) upon treatment with MMV665953 (Figure 2C). We therefore attempted to locate where this activity was occurring within the typical ER-secretion pathway using the MSP1-Nluc cell line wherein the Nluc reporter construct is directed to the PV but not trafficked into the iRBC (Figure 1D). These assays were completed with MMV396797 and MMV665953 only, as MMV666687 was not commercially available. Following drug treatment, secretion of MSP1-Nluc was analysed using our export assay (Figure 3C), showing that treatment with Brefeldin A resulted in retention of MSP1-Nluc signal in the parasite compared to the DMSO control (unpaired t test, p = 0.0001). After treatment with Torin 2 and MMV396797, MSP1-Nluc was largely located in the PV, however both displayed significantly increased signal retained in the parasite when compared to DMSO treated cells (unpaired t test, Torin 2 p = 0.0026, MMV396797 p = 0.0023). In parasites treated with MMV665953 MSP1-Nluc was blocked almost entirely within the parasite (85.2 ± 0.1 % retained signal). This indicated that inhibitors that block export of a PEXEL containing Hyp1-Nluc reporter (Figure 3C) also act upon the PV resident MSP1-Nluc reporter suggesting shared elements between the trafficking pathways.

### Titration of export inhibitor reveals potency against export pathway

To determine the compound concentrations required to optimally block export, we completed nanoluciferase export assays at a range of drug concentrations. First for comparison, export assays were performed using parasites treated with DMSO (at 0.1% and 1% to cover the range of DMSO concentrations administered with MMV compounds) with known export inhibitors Brefeldin A (5 µg/mL) and Torin 2 (10 nM) acting as positive controls (Figure 4A). DMSO had a minimal effect upon export even when used at 1% v/v for 5 hours. For MMV396797 and MMV665953, concentrations were progressively halved from a maximum concentration of 50 μM to a minimum of 0.78 μM (Figure 4B). Reporter protein was retained in both the parasite and PV compartments following treatment with MMV396797 (Figure 4B). These results are consistent with a target for MMV396797 in the secretory/export pathway, at or upstream of PTEX trafficking across the PVM. Interestingly, the export inhibiting activity of MMV396797 reached a plateau at around 12.5 µM resulting in a maximum retained/secreted fraction of 54.1 ± 2.9 % (export EC_50_ = 0.94 µM, curve capped at maximum observed export inhibition). Similarly, the decrease in overall cellular Hyp1-Nluc signal also plateaued around this concentration (Figure 4C). As Brefeldin A and Torin 2 controls appear to inhibit export to a similar level as high concentrations of MMV396797, it is possible that this is the maximum level of true export inhibition measurable by this assay after accounting for protein exported before drug treatment. Alternatively, it is possible that MMV396797 is not able to fully block the export pathway and the plateau of total nanoluciferase signal observed may represent the maximal effect MMV396797 can have on the parasite within the 5 hour treatment window. In contrast, continually increased retention of Hyp1-Nluc in the parasite was observed in a dose-dependent manner with increasing concentrations of MMV665953 (export EC_50_ = 10 µM, curve capped at maximum observed export inhibition). The total cellular Hyp1-Nluc signal was dramatically reduced with increasing concentration of MMV665953, likely indicating a decrease in parasite viability that appears to plateau at 25 µM (Figure 4C). It is possible that at high concentrations MMV665953 strongly inhibit parasite growth causing knock-on effects that impact export, and that the resulting low levels of nanoluciferase signal may accentuate the protein retention readout of the assay.

**Figure 4.**
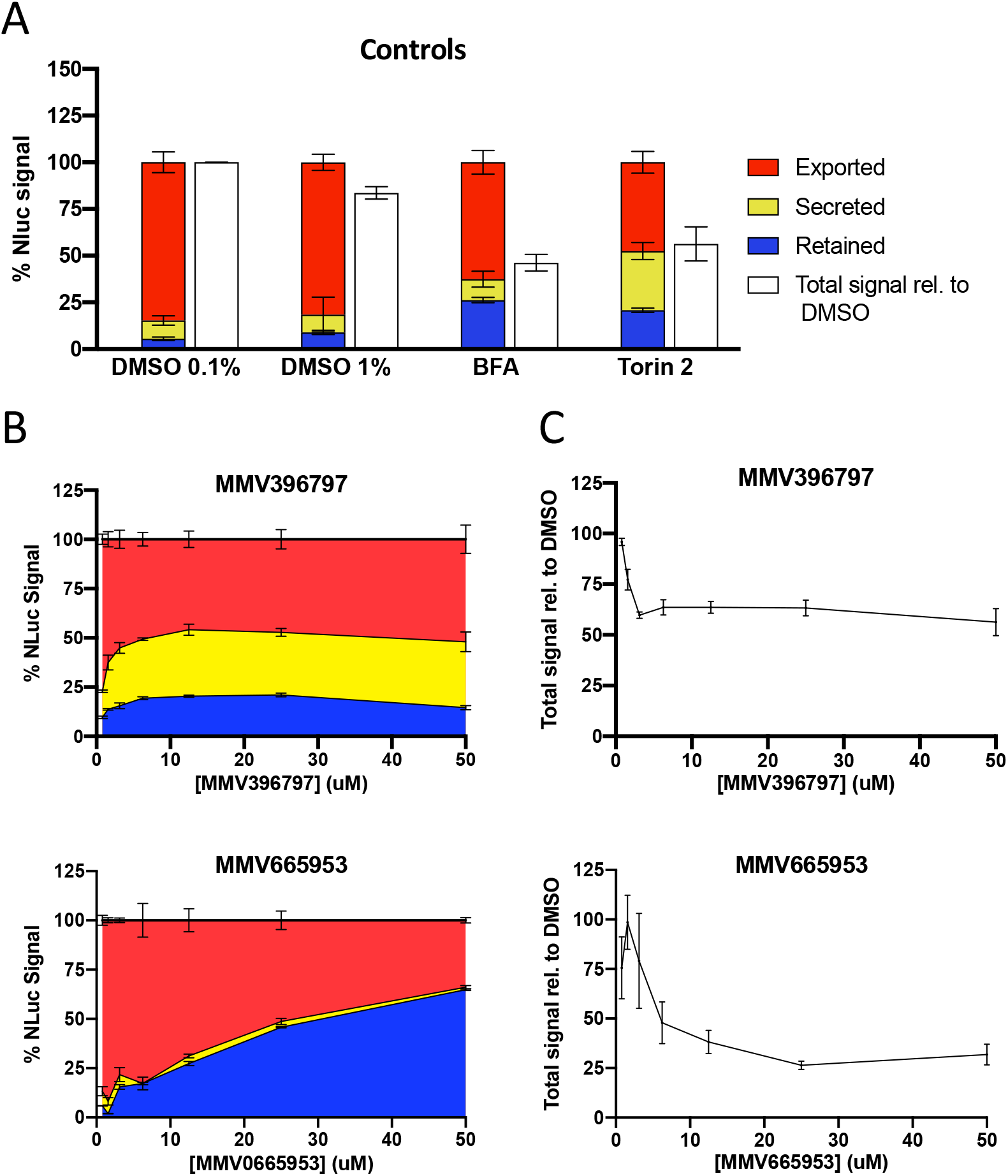
Export inhibitors trap exported proteins in a dose-dependent manner. **A)** Export assays showing the effect of control compounds and several DMSO concentrations. Synchronous 20-24 hpi Hyp1-Nluc parasites were treated for 5 hours with DMSO (0.1%), DMSO (1%), Brefeldin A (5 μg/mL) or Torin 2 (10 nM). Export assays were performed as described in Figure 1A and exported (red), secreted (yellow) retained (blue) and total (white) nanoluciferase signal was plotted. **B)** Export assay analysis of a range of export inhibitor concentrations. Synchronous (20-24 hpi) parasites expressing Hyp1-Nluc were treated with MMV396797 and MMV665953 at a range of concentrations decreasing from 50 μM to 0.78 μM in 2-step dilutions for 5 hours before export assays were performed. **C)** Total cellular nanoluciferase activity as measured from the experiments presented in B), normalised to total signal from the DMSO (0.1%) control group from A). Experiments were performed in technical duplicates over 3 biological replicates. All retained, secreted, exported and total nanoluciferase values in A-C) are presented as mean percentage ± SEM.

### Recovery of protein export following removal of inhibitors

To determine the reversibility of the export-inhibiting activity of MMV396797 and MMV665953, we treated parasites with either compound for 3 hours before allowing them to recover for 3 hours following drug treatment before measuring protein export at both time points. Some of the 3 hour parasites were also treated with drugs for an additional 3 hours (total treatment of 6 hours). Parasites allowed to recover for 3 hours after treatment with brefeldin A displayed significantly increased export of Hyp1-Nluc signal compared to parasites that were not allowed to recover (Figure 5A-B, 3 hour treatment vs 3 hour recovery) (unpaired t-test, p = 0.0033). In contrast, Hyp1-Nluc export in Torin 2 treated parasites only slightly recovered within the 3 hour recovery window after removal of drug (unpaired t-test, ns p = 0.052), and the amount of reporter trapped within the parasite continued to increase with prolonged drug treatment (6hr treatment) indicating that the parasite was still producing protein but it was unable to be exported (Figure 5C). Interestingly, after removal of MMV396797 for 3 hours, nanoluciferase signal detected in the RBC space was significantly increased (unpaired t-test, p = 0.017), showing recovery of protein export following removal of the drug (Figure 5D). Some signal remained in the PV and parasite fractions after drug removal, suggesting that not all protein blocked by the export inhibitor was able to be exported within the 3-hour time frame (Figure 5D). After removal of each drug, total Hyp1-Nluc signal moderately increased compared to a 6-hour treatment however these increases were not significant (unpaired t-test p > 0.05). It is possible that lowered nanoluciferase signal results from reduced parasite viability in combination with degradation of Nluc reporter protein blocked in the export pathway. Removal of drug would therefore only partially restore signal as the reporter continued to be expressed and exported in viable parasites. Removal of MMV665953 resulted in a dramatic increase in exported nanoluciferase signal (unpaired t-test, p = 0.0003) but no recovery of severely lowered total nanoluciferase signal (unpaired t-test p > 0.05) (Figure 5E). As such it seems likely that MMV665953 has a strong and rapid impact on parasite viability and that the observed recovery of protein export is a side effect magnified by the low total nanoluciferase signal (consistent with Figure 4C).

**Figure 5.**
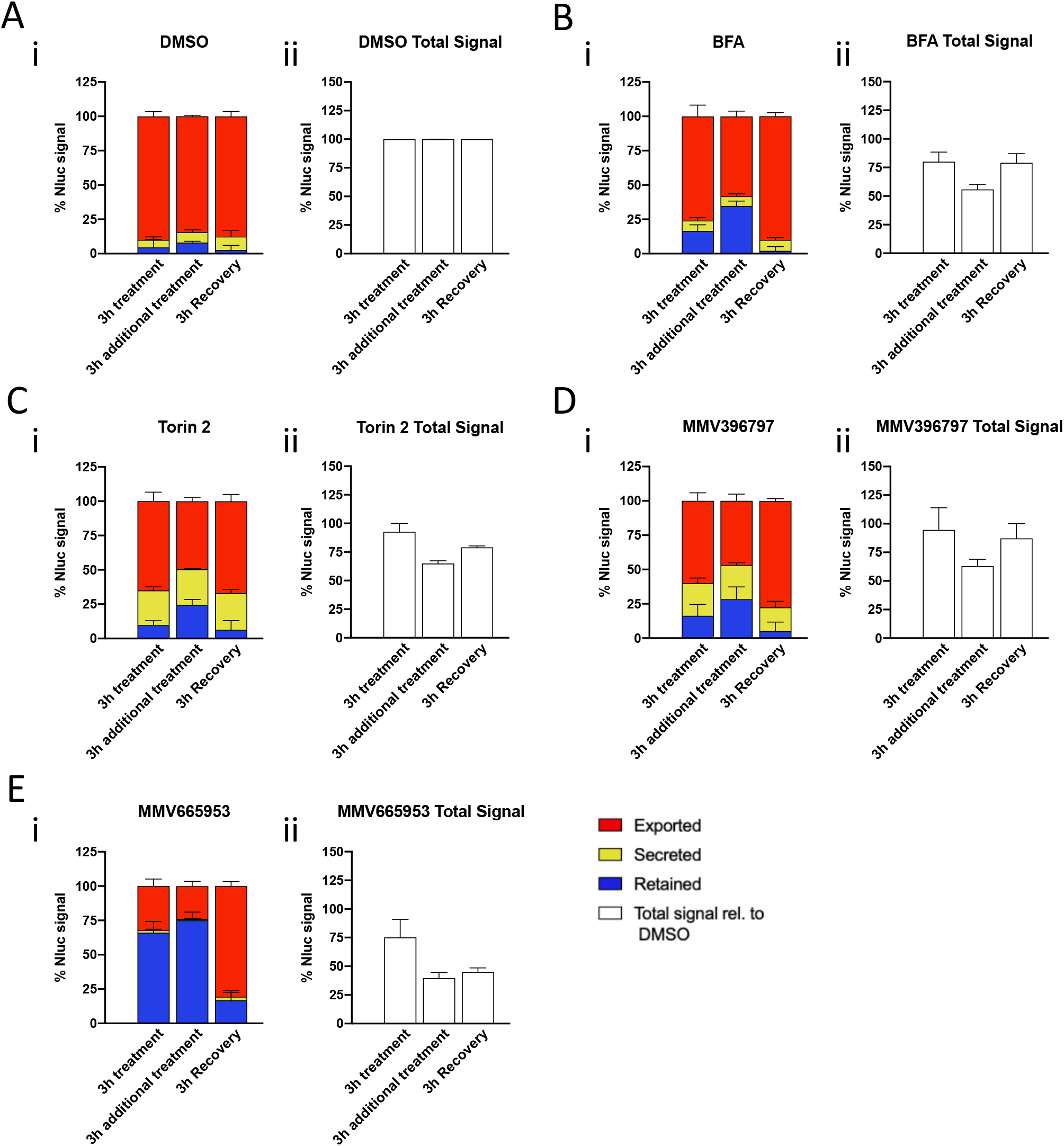
Recovery of protein trafficking after removal of export inhibitors. Export assays showing recovery of export after removal of export inhibitors. Hyp1-NLuc parasites were synchronised to 20-24 hpi and treated with **A)** DMSO (0.1 %), **B)** Brefeldin A (5 μg/mL), **C)** Torin 2 (10 nM), **D)** MMV396797 (4.8 μM) or **E)** MMV665953 (20.6 μM). **A-Ei)** After 3 hours a group was selectively lysed with equinatoxin II, saponin or Igepal-630 to release exported (red), secreted (yellow) and retained (blue) nanoluciferase-reporter. The remaining cells were grown for a further 3 hours in 2 groups, either with drug (3h additional treatment) or after removal of drug (3h recovery) before selective lysis and measurement by export assay. **A-Eii)** Total cellular nanoluciferase activity was measured for each treatment group in part i) and normalised to the signal from the DMSO vehicle control. Experiments were performed in technical duplicates over 3 biological replicates. All retained, secreted, exported and total nanoluciferase values in A-E) are presented as mean percentage ± SEM.

### Immunofluorescence assays reveal MMV396797 traps cargo within the parasite and parasitophorous vacuole

We next investigated the export-blocking characteristics of MMV396797 and MMV665953 by visualising the location of trapped Hyp1-Nluc cargo by immunofluorescence labelling and widefield microscopy. To assess the location of Hyp1-Nluc, primary antibodies against the PVM protein EXP2 ^43^ and against nanoluciferase ^65^ were used to label the PVM and exported fusion protein respectively. In trophozoite stage parasites (aged 25-29 hpi) treated with DMSO for 5 hours, exported Hyp1-Nluc reporter can be seen within the boundary of the host RBC, and what is likely newly synthesised Hyp1-Nluc protein is present within the parasite compartment (Figure 6A). Following 5-hour treatment with Brefeldin A or Torin-2, the Hyp1-Nluc reporter was observed within the bounds of the PVM and appeared to strongly concentrate within the parasite (Figure 6A). Following treatment with MMV396797, Hyp1-Nluc signal was also concentrated within the parasite (Figure 6A). We note that in the export assays for Torin 2 and MMV396797 (Figures 2A,C & 4 A,B) there was both an increase in the proportion of Hyp1-Nluc signal in the parasite and PV but by microscopy we did not observe a matching of the increase in PV signal which would have colocalised with EXP2. This will be discussed in a later section. Treatment with MMV665953, resulted in reduction of Hyp1-Nluc within both the RBC space as well as the parasite (Figure 6A).

**Figure 6.**
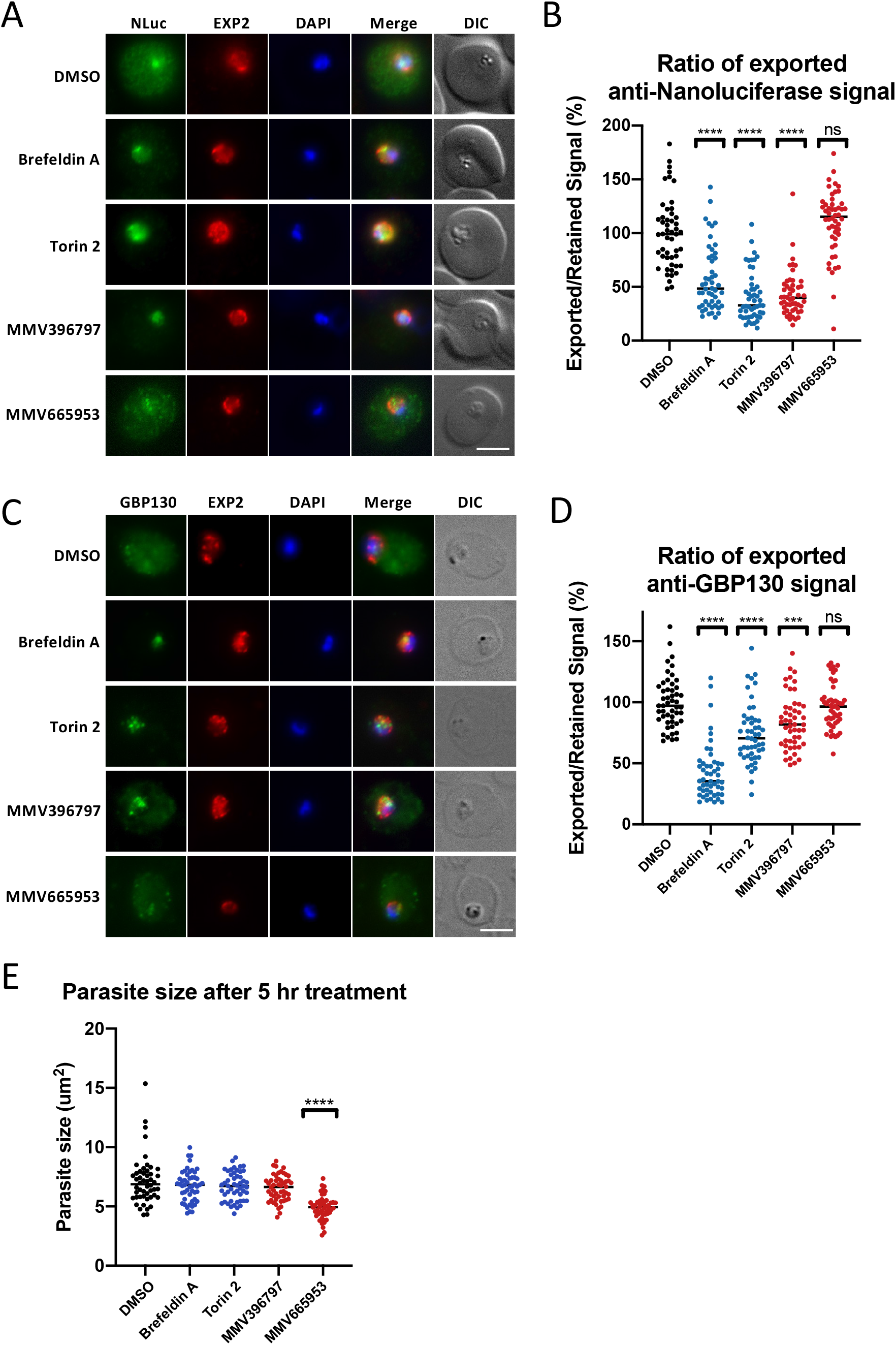
Widefield immunofluorescence microscopy shows retention of exported proteins within the bounds of the PVM. **A)** Widefield fluorescence microscopy images of Hyp1-Nluc parasites treated with DMSO (0.1 %), Brefeldin A (5 μg/mL), Torin 2 (10 nM), MMV396797 (4.8 μM) or MMV665953 (20.6 μM) for 5 hours from 20-24 hpi. Labelling was performed with mouse anti-EXP2 and rabbit anti-nanoluciferase primary antibodies and goat anti-mouse Alexafluor 594 and goat anti-rabbit Alexafluor 488 secondary antibodies. Scale bar = 4 μm. **B**) The ratio of anti-Nluc signal within the iRBC and PV measured from cells imaged in A). The perimeter of the iRBC and PV were manually traced and the anti-Nluc signal within each compartment was measured using ImageJ. Exported signal was divided by signal in the PV to produce an export ratio before being normalised to DMSO. **C)** Widefield fluorescence images of Hyp1-Nluc parasites treated as specified in A) from 13-15 hpi. Cells were labelled with rabbit anti-EXP2 and mouse anti-GBP130 (WEHI antibody facility) primary antibodies and anti-rabbit Alexafluor 594 and anti-mouse Alexafluor 488 secondary antibodies. Scale bar = 4 μm. **D)** The ratio of anti-GBP130 signal within the iRBC and PV measured from cells imaged in C). GBP130 signal was measured and an export ratio was calculated as described in B). **E)** Parasite size was measured by manually tracing the perimeter of the PV and measuring the area within this space using ImageJ. The PV was visualised using anti-EXP2 signal as a marker. For panels B,D,E) The statistical difference of the mean of each treatment compared to DMSO was calculated using an unpaired t-test (* p < 0.05, ** p ≤ 0.01, *** p ≤ 0.001, **** p ≤ 0.0001). n = 50 cells for each treatment group.

Following the approach to quantify export inhibition used for plasmespin V blocking compounds ^40^, data from our immunofluorescence assays was quantitatively analysed to examine the ratio of exported to retained signal across a wider population of parasites. Signal within the bounds of the PVM and outside the PVM but within the RBC membrane was measured over 50 cells for each drug treated parasite population. The ratio of exported and PV-contained signal was used to relatively quantify export inhibition. When compared to DMSO-treated cells, Brefeldin A, Torin 2 and MMV396797 treatment resulted in significantly reduced export of Hyp1-Nluc (unpaired t-test, Brefeldin A, Torin 2 and MMV396797 p < 0.0001)(Figure 6B). Treatment with MMV665953 did not significantly reduce export of Hyp1-Nluc relative to that present within the parasite (unpaired t-test, p = 0.11).

We next sought to investigate the effect of our export inhibitors on the trafficking of an endogenously expressed exported protein (glycophorin binding protein 130 (GBP130))^66^. Hyp1-Nluc parasites were drug treated as described above and imaged at 18-20 hpi. Consistent with previous data, in control parasites treated with DMSO for 5 hours, GBP130 was observed to be exported into the RBC compartment with some protein present within the parasite (Figure 6C). In parasites treated with Brefeldin A, Torin 2 and MMV396797, the majority of the GBP130 signal was retained within the parasite relative to the RBC compartment. In comparison, MMV665953 treatment greatly reduced the anti-GBP130 signal within the parasite in contrast to a modest signal in the RBC space (Figure 6C). This could in part be due to the broader expression window of GBP130 compared to the Hyp1-Nluc reporter making it likely that some GBP130 was exported before drug treatment commenced. Quantification of signal over 50 cells showed that GBP130 was significantly retained within the PV after treatment with Brefeldin A, Torin 2 and MMV396797 (unpaired t-test, Brefeldin A and Torin 2 p < 0.0001, MMV396797 p = 0.0003) but not with MMV665953 (unpaired t-test, p = 0.42)(Figure 6D).

This finding was confirmed in a second cell line expressing a hemagglutinin epitope-tagged lysine-rich membrane-associated PHIST protein (LyMP) which is exported to the infected RBC membrane in untreated parasites ^67^. Drug treatment for 5 hours from 20-24 hpi and labelling with anti-HA antibodies revealed that LyMP-HA was visually retained in the same manner as GBP130 and Hyp1-Nluc for all compounds except MMV665953 (Figure S2).

A possible confounding factor for the data presented above is the size of the parasite. Specifically, the size of *P. falciparum* parasites increases as they progress through the asexual cycle, and the timing of protein export also varies between proteins ie. some proteins begin to be exported earlier (GBP130) while others are exported later (Hyp1-Nluc). Inhibition of parasite growth could therefore not only affect the size of the parasite, but also protein export. To determine whether inhibition of parasite growth was contributing to the observed retention of exported proteins, we measured the relative size of parasites after drug treatment. The perimeter of the PV membrane was measured from 50 cells that were previously labelled and imaged (Figure 6A). Cells treated with Brefeldin A, Torin 2 or MMV396797 were not significantly different in size to those treated with DMSO (Figure 6E) (unpaired t-test, p > 0.05). Parasites treated with MMV665953 were significantly smaller than DMSO-treated counterparts, suggesting that this compound strongly inhibits parasite growth (unpaired t-test, p < 0.0001). With MMV396797 emerging as the most specific inhibitor of protein export in the MMV Malaria Box we sought to find its protein target through selection of parasites resistant to the compound. Point mutations or copy number variants in the gene for the drug’s target protein could then be identified through whole genome sequencing. Several attempts to select for parasites resistant to cyclic treatment with 10x EC_50_ of MMV396797 produced surviving parasites but subsequent growth assays indicated they had not become resistant to MMV396797 relative to the original parental population (Figure S3).

### Inhibition of protein export impairs iRBC rigidity

Exported proteins perform a wide range of functions in the RBC, many of which are vital to parasite survival and virulence ^68,69^. To further elucidate the export-inhibiting properties of our lead compounds, we assessed several phenotypes modulated by exported proteins downstream of the export pathway. As the asexual-stage parasite develops it progressively rigidifies its host RBC which can largely be attributed to the binding of exported proteins to the RBC membrane skeleton ^51,70^. We therefore sought to measure the relative rigidity of infected RBCs after treatment with export inhibitors. To achieve this, infected RBCs were synchronised to a 14-16 hpi window and drug treated for 5 hours for completion of rigidity assays at 19-21 hpi. This time window was selected to pre-empt the rigidification of early trophozoite iRBCs by exported proteins that interact with the iRBC membrane skeleton ^51,71^. Infected RBCs were flowed through a bed of microbeads such that highly rigid cells were caught in the microbead matrix, while more deformable cells were eluted (Figure 7A) ^72^. Assessment of the input and eluted fractions by Giemsa stained, dried blood smears showed that compared to a DMSO vehicle control, Brefeldin A and MMV396797 treated cells both showed significantly reduced rigidity (unpaired t-test, Brefeldin A p = 0.0009, MMV396797 p = 0.0026), likely due to export inhibition (Figure 7B). Treatment with MMV665953 at 14-16 hpi resulted in severely growth-inhibited parasites, thus they could not be assessed by microfiltration (Figure S4).

**Figure 7.**
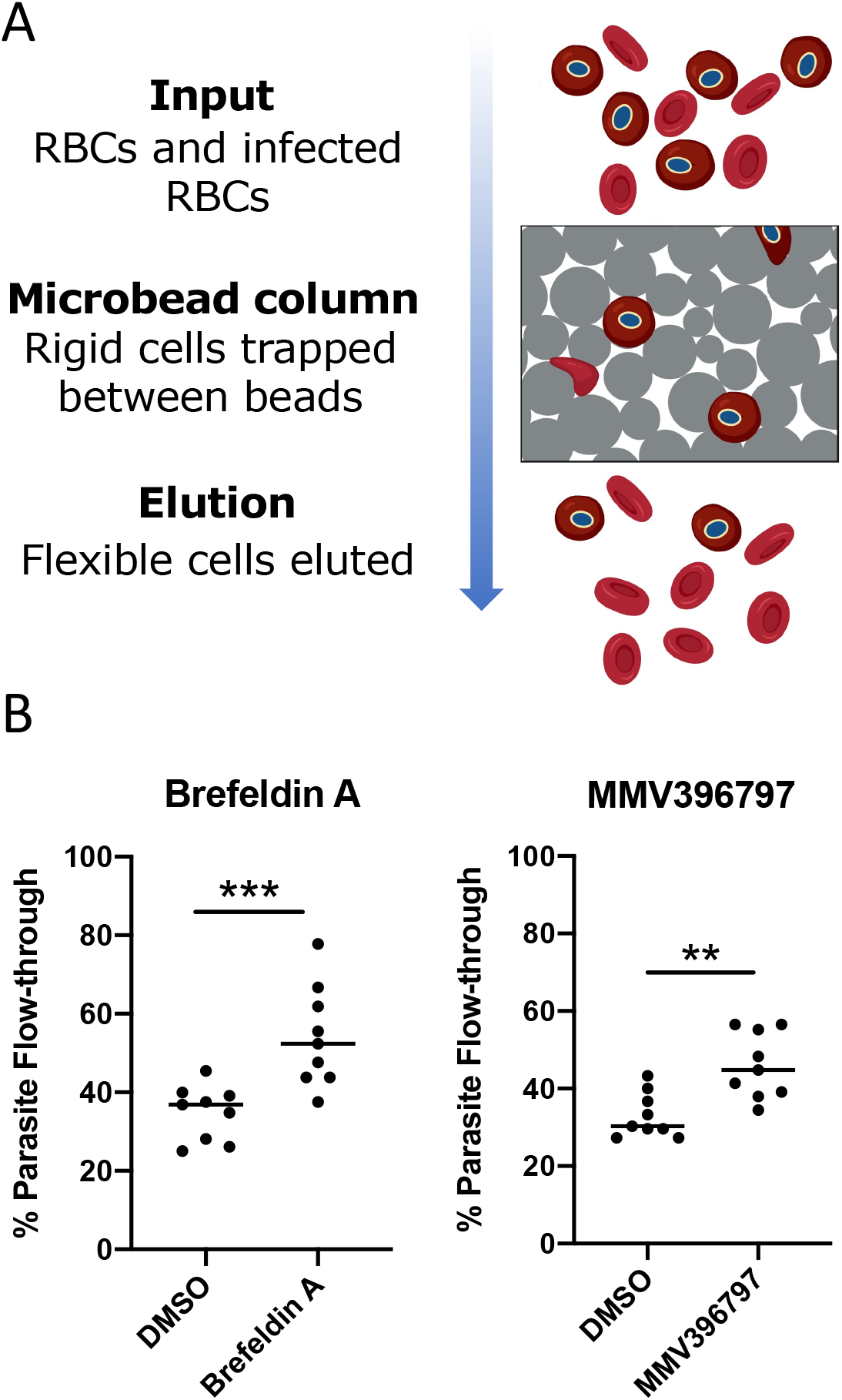
Inhibition of protein export impairs host-cell rigidity. **A)** Diagrammatic representation of measurement of relative cellular rigidity by microfiltration. iRBCs were passed through a bed of microbeads. Relatively rigid cells were trapped in the microbead matrix while deformable cells were eluted. The parasitaemia of the input and eluted fractions was analysed using Giemsa stained, dried blood smears and the parasite flow-through was calculated as a percentage. **B)** Relative rigidity of export inhibitor-treated iRBCs. MSP1-NLuc parasites were synchronised to 14-16 hpi and treated with Brefeldin A (5 μg/mL) or MMV396797 (4.8 μM) for 5 hours. A duplicate culture treated with DMSO as a vehicle control was also prepared for each experiment. The relative rigidity of cells was measured by microfiltration as described in A). Experiments were performed in technical triplicates over 3 biological replicates. All flow-through values are presented as mean percentage ± standard deviation. The statistical difference of the mean of each treatment compared to DMSO treatment groups was calculated using a nonparametric unpaired t-test. ** p = 0.0026, *** p = 0.0009.

### MMV396797 reduces infected RBC binding to an endothelial ligand under flow conditions

To avoid becoming trapped and subsequently destroyed in the spleen, *P. falciparum* parasites mediate cytoadherence of their host RBC to blood vessel endothelial walls thus removing the cell from circulation. This requires the presentation of exported PfEMP1 proteins through the RBC membrane with several other exported parasite proteins acting as stabilisers or RBC remodelling agents ^73–75^. Inhibition of protein export would prevent arrival of these proteins at the RBC surface, thus leaving the parasite vulnerable to human defences. We analysed cytoadherence of export inhibitor-treated iRBCs against the endothelial ligand chondroitin sulfate-A (CSA) under physiological flow conditions. CSA is a glycosaminoglycan to which binding is prevalent in cases of placental malaria ^76^. Parasite cytoadhesion in the placenta is mediated by the PfEMP1 variant VAR2CSA which is specifically responsible for CSA binding ^77,78^. Following a 5-hour drug treatment, CS2 parasites (expressing VAR2CSA) at 20-25 hpi were passaged through a CSA-lined microchannel (Figure 8A). The number of cells that bound to the channel was analysed by widefield microscopy of several randomly imaged fields of view (Figure 8B). Compared to those treated with DMSO, iRBCs treated with Brefeldin A or MMV396797 bound in significantly lower numbers per area, at 0.66 ± 0.04 and 0.65 ± 0.04 times the binding density respectively (unpaired t-test, p < 0.0054) (Figure 8C). To account for cells binding non-specifically to the channel, a control microchannel was prepared without CSA. Cells bound to these channels at a density of 0.004 ± 0.002 times that of CSA-coated chambers, indicating minimal non-specific binding.

**Figure 8.**
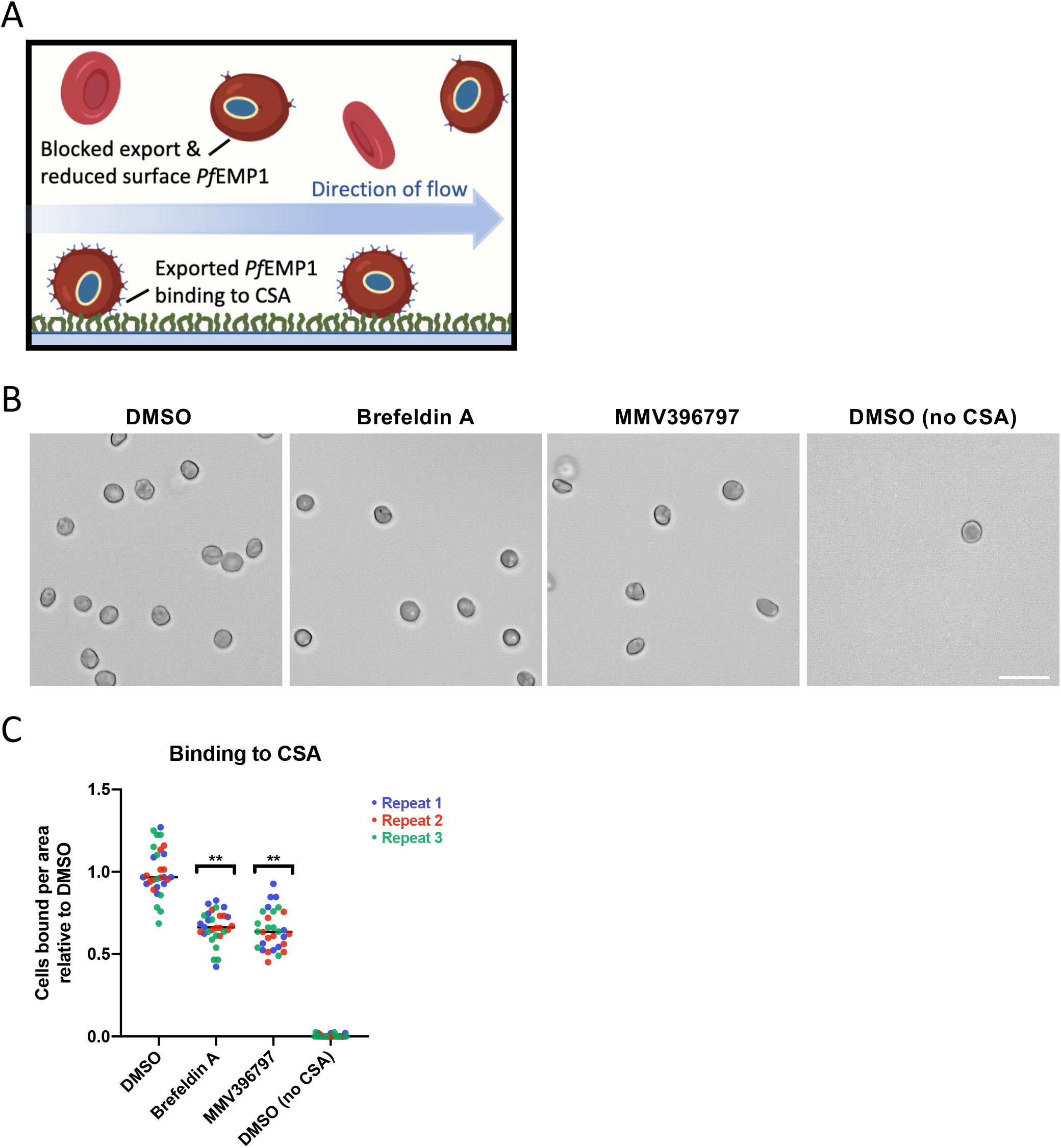
Reduced binding to endothelial ligands following treatment with export inhibitors. **A)** Diagram depicting iRBC binding to CSA under flow conditions. iRBCs were flowed through a CSA-coated microchannel after treatment with export inhibitors. Cells presenting *Pf*EMP1 (VAR2CSA) at the RBC surface adhere to the CSA coating, while cells with less *Pf*EMP1 at the RBC surface (as a result of export inhibition) are less likely to successfully adhere. **B)** Representative widefield light microscopy images of CS2-infected RBCs bound to a microchannel under flow conditions as described in A). Before being passed through the channel parasites were synchronised to 15-19 hpi and treated with DMSO (0.1 %), Brefeldin A (5 μg/mL), or MMV396797 (4.8 μM) for 5 hours. DMSO, Brefeldin A and MMV396797-treated cells were passed through CSA-lined channels, while a second DMSO treatment group was passed through a control channel with no CSA coating. Scale bar = 20 μm. **C)** CS2 parasites were synchronised, drug treated and flowed through microchannels as detailed in B). Bound cells were counted over 10 random regions (each 281.26 x 178.14 μm) along the length of the channel for each of 3 biological repeats. The number of bound cells per area was presented after normalisation against the DMSO treatment group. The statistical difference of the mean of each treatment compared to the DMSO treatment group flowed through a CSA-lined channel was calculated using an unpaired t-test. ** Brefeldin A p = 0.0053, MMV396797 p = 0.005.

## Discussion

To find novel inhibitors that could rapidly reduce harm to the human host whilst killing *P. falciparum* blood stage parasites we screened the MMV Malaria Box for compounds that could inhibit parasite protein secretion and export. From this screen, 14 compounds were found that appeared to reduce the bioluminescence of the Hyp1-Nluc reporter protein in the exported RBC compartment relative to the parasite and PV compartments. Subsequent counter screening indicated 10 of these compounds were false positives as they inhibited nanoluciferase activity, 7 to a lesser extent in a detergent buffer, artefactually inflating the parasite signal compared to the exported reporter signal. Only three compounds MMV396797, MMV665953 and MMV666687, did not show this bias and were selected for cell-based assays to confirm they blocked protein export and phenomena dependent on protein export. Unfortunately, MMV666687 could not be commercially obtained at the time of this work and so no further characterisation was performed. Of the other two remaining compounds, only MMV396797 was shown to robustly block protein export by microscopy but its protein target could not be identified through the recovery of drug resistant parasites. MMV396797 has a chemical structure that belongs to the pyrazolopyrimidine class and this chemotype shares distinct similarities to kinase-like inhibitors. Chemogenomic and metabolic profiling implies that the target of MMV396797 is involved in haemoglobin catabolism which interestingly matches the signature of the PI4K inhibitors, Torin 2 and MMV390048 ^79,80^. It is possible that inhibition of vesicular trafficking mediated by PI4K inhibitors could reduce the delivery of haemoglobin containing vesicles to the food vacuole for digestion thereby reducing haemoglobin catabolism.

It was noted that the mechanism of action of MMV396797 was more Torin 2-like with reporter protein trapped in both the parasite and PV rather than Brefeldin A-like in which the reporter tended to be mostly trapped in the parasite. Brefeldin A acts early in the secretory pathway by inhibiting vesicular transport between the Golgi and the ER leading to the collapse of these organelles into the one body ^81^. Brefeldin A is unsuitable as an anti-malarial starting point as it is active against many eukaryotes and therefore toxic to the host. Torin 2 has been shown to be highly potent against blood stage parasite growth and to be effective against multiple parasite stages ^28^. Torin 2 was shown to interrupt protein trafficking but attempts to select for resistance were unsuccessful and the compound’s protein target and mechanism of action could not be fully derived. Recently, derivatives of Torin 2 that have improved selectivity for parasites and better solubility and microsomal stability have been developed ^27^. Parasite resistance to one of these compounds, NCATS-SM3710, has been selected and the target is *Pf* PI4KIIIβ which phosphorylates phosphatidyl inositol (PI) to produce phosphatidyl inositol 4 phosphate which is important for signalling and lipid trafficking between the Golgi, trans-Golgi and plasma membrane of the parasite. NCATS-SM3710 and Torin 2 do not appear to be structurally related to MMV396797 but the similarity with which they trap reporter cargo in both parasite and PV suggest their protein targets could function within the same pathway.

MMV665953 is closely related to MMV665852 and whole-genome sequencing of resistant MMV665852 clones, uncovered PfATPase2 as a possible target of the diaryl urea class ^82,83^. MMV665852 was also identified as an inhibitor of our export assay but excluded in our counter screen due to inhibition of the Hyp1-Nluc reporter. PfATPase2 is essential for parasite survival and is likely involved in the maintenance of lipid asymmetry in the bilayer which is important for membrane trafficking ^84^. It is possible that MMV665853’s inhibition of PfATPase2 so strongly reduced parasite growth, that by microscopy it did not appear to convincingly reduce protein export as much as the other inhibitors. PfATPase2 is upregulated by phosphatidylinositol 4-phosphate ^84^ the product of PI4KIII*β* which is inhibited by Torin 2 and so we note the link between components of the vesicular trafficking system.

As mentioned above, MMV396797 produced an effect more like Torin 2 rather than Brefeldin A with MMV396797 and Torin 2 trapping the reporter in both the parasite and PV rather than only in the parasite as Brefeldin A did. By immunofluorescence microscopy however, we did not observe the PV trapping effect for MMV396797 and Torin 2, and instead observed almost complete trapping of Hyp1-Nluc inside the parasite as was also the case for native GBP130 and LyMP proteins. A possible reason for this discrepancy, is that Torin 2’s inhibition of PfPI4KIII*β* between the Golgi and parasite plasma membrane could lead to some leakiness following saponin treatment allowing cytoplasmic proteins to escape thereby increasing the PV fraction in the export assay but not by microscopy. On the other hand, Brefeldin A which acts upstream between the ER and Golgi may better trap cargo in the parasite fraction following saponin treatment in agreement with the western blot data showing that ER protein ERC, was highly resistant to leaking into the PV following saponin treatment.

Interestingly, of the 10 export inhibitors removed by the counter screen, six have previously been identified as PfATP4 inhibitors (MMV000642, MMV000662, MMV006429 MMV665803, MMV006764 and MMV006656)^22^. PfATP4 is an ATP dependent sodium efflux pump that exchanges one Na^+^ ion for a proton and its inhibition causes an increase in both Na^+^ levels and pH within the parasite. This causes the parasites to swell, and dysregulation of cellular homeostasis rapidly inhibits parasite growth ^22,85^ which may have indirectly decreased protein export. Why PfATP4 inhibitors should also differentially influence Hyp1-Nluc reporter activity +/- detergent in the counter screen is not known.

An essential component of the protein export pathway is PTEX (Plasmodium Translocon of Exported proteins) and given its essentiality ^37,39^ we had hoped to identify specific PTEX inhibitors. PTEX is located at the PVM where it unfolds and extrudes protein cargo across the PVM into the RBC compartment ^36,37,39,44^. Knockdown of HSP101 and the other core component proteins of PTEX (PTEX150 and EXP2) often cause PEXEL proteins to accumulate as large bodies within the PVM which are clearly discernible by immunofluorescence microscopy ^37,39,65,86^. Furthermore, trapping PEXEL reporters rendered unfoldable through chemical stabilisation of a murine dihydrofolate dehydrogenase domain within PTEX caused extensive morphological changes at the PVM ^65^. Although the Hyp1-Nluc export assay indicated MMV396797 caused increased trapping of the reporter in the PV, immunofluorescence microscopy of MMV396797-treated parasites indicated the PEXEL reporter and endogenous exported proteins were primarily trapped inside the parasite and did not resemble proteins trapped at the PV. As MMV396797 appeared by microscopy to block protein export within the parasite, it seems unlikely that this compound targets PTEX150 or EXP2, which reside at the PVM. While HSP101 is dually located at the PVM and the parasite ER ^46^, the relative lack of PEXEL protein blocked in the PV space after MMV396797 treatment suggests that it is unlikely to also be the target of this compound.

With respect to rigidification of the iRBC, MMV396797 acted like Brefeldin A in blocking a process which is largely caused by exported parasite proteins binding and remodelling the iRBC membrane skeleton. One of the most well studied proteins that contributes to rigidification is the PEXEL protein KAHRP, which binds to the RBC membrane skeleton producing raised knob-like protrusions that project PfEMP1 proteins out from the iRBC surface to promote cytoadherence ^71,75^. Knobs structurally link PfEMP1 to the iRBC membrane skeleton, thus it is possible that they also promote cytoadherence by using the newly rigidified skeleton as a scaffold to brace against the forces of binding under flow conditions ^87^. Although we were unable to directly determine MMV396797’s effect on KAHRP trafficking, the compound’s inhibition of export of the iRBC cytoskeleton binding protein, LyMP, and the iRBC’s lack of rigidity strongly indicated that export of KAHRP into the iRBC was blocked. Similarly, the lack of cytoadherence of CS2 iRBCs to CSA coated surfaces likewise indicated there was a lack of export of VAR2CSA, KAHRP and/or other proteins involved in rigidification. Treatment of ring stage parasites with MMV396797, before but not after they have exported rigidification and cytoadherence factors could therefore reduce the capacity of the iRBC to do harm while the compound kills the parasite due to dysregulation of protein trafficking.

Thus far, we have only investigated the roles of secretion and export inhibitors during the early trophozoite stage, but it is anticipated these inhibitors could also block RBC invasion due to the reduction of protein transport to rhoptry and microneme invasion organelles and post-invasion, dense granules. It is noted that predicted PI4K inhibitors in the MMV Pathogen Box, MMV020391 and MMV085499, reduced invasion after only four hours treatment of schizonts ^24,29,88,89^ therefore indicating that reduction of protein trafficking is highly deleterious for invasion. The ability of Malaria Box compounds to inhibit schizont to ring transition in *P. falciparum* has been previously screened to identify novel invasion inhibitors and MMV396797 was flagged as a relatively effective inhibitor ^25^. Additionally, MMV396797 appears to be relatively potent against gametocytes and liver-stage parasites ^19,23, 90–92^ suggesting the strategy of targeting protein trafficking could block the parasite’s lifecycle at multiple stages thereby satisfying several target candidate profiles which novel antimalarial compounds should ideally have ^93^.

## Conclusions

Here we have used a novel screen to find several inhibitors of protein secretion and export in the MMV Malaria Box. Follow up counter screens and fluorescence microscopy indicated MMV396797 was the most specific protein export inhibitor, highlighting the need to use independent cell-based assays to triage primary screen hits. Our results also showed that MMV396797 impaired iRBC rigidity and cytoadherence and may therefore simultaneously kill the parasite and reduce its virulence. To define the mechanism of action of MMV396797, we attempted to select for drug resistant parasites to identify the compound’s protein target but were unsuccessful. This could indicate that MMV396797 may be an ‘irresistible’ compound and other methods might be required such as chemo-proteomic approaches to identify parasite proteins that are the biological target of the compound.

## Materials & Methods

### Parasite lines, culture and synchronisation

The *Plasmodium falciparum* parasite lines used in this study are as follows:

3D7, CS2 ^94^, 3D7-Hyp1-nanoluciferase ^58^, 3D7 MSP1-nanoluciferase ^58^, 3D7 PV1-HA-glmS^59^ and 3D7 LyMP-HA ^95^.

*Plasmodium falciparum* parasites were cultured in human RBCs (Australian Red Cross Blood Bank, Type O+) at 4% haematocrit in NaHCO_3_ and AlbumaxII/human serum supplemented RPMI (RPMI (Sigma Aldrich), 25 mM HEPES (GIBCO), 25 mM NaHCO_3_ (Thermo Scientific), 365 μM hypoxanthine (Sigma Aldrich), 31.25 μg/mL of Gentamicin (GIBCO) and 0.5% AlbumaxII (GIBCO) or 0.25% AlbumaxII and 5% v/v heat-inactivated human serum (Australian Red Cross)) at 37°C. Cultures were grown under a low-oxygen gas mix (94% N_2_, 5% CO_2_, 1% O_2_). Transfectant parasites used in this study expressing human dihydrofolate reductase (hDHFR) were selected and maintained in culture with WR99210 (WR, Jacobus Pharmaceutical Company) at 2.5 nM. Parasite age and parasitaemia was monitored by brightfield microscopy of blood slides fixed in 100% methanol (Thermo Fisher) and stained for 10 minutes using 10% (v/v) Giemsa (Merck).

To select for knob-positive parasites, cultures were periodically enriched by gelatin floatation^96^. Briefly, infected RBCs were collected from culture by centrifugation (500 g, 5 minutes) and resuspended in 0.75% (w/v) gelatin (DIFCO) in RPMI. Resuspended cells were allowed to separate into a pellet and supernatant fraction by incubation at 37°C for 45 minutes. The supernatant fraction was collected by centrifugation (500 g, 5 minutes) and returned to culture.

For use in experiments, parasites were synchronised to a 2–4-hour time window. Schizont-infected RBCs were enriched by a 65% (v/v) Percoll gradient ^97^(Sigma Aldrich) from 30 mL cultures at 5-10% parasitaemia. The collected schizonts were returned to culture in 200 μL RBC and 5 mL complete culture medium incubated at 37°C shaking at 60 RPM for 2-4 hours. After incubation newly invaded infected RBCs were selected for by incubation in 5% sorbitol (Sigma Aldrich) for 10 minutes at 37°C. Synchronous parasites were returned to culture for later use.

### Compounds

The 400 Malaria Box compounds used for the export screen and counter screen were provided by MMV and were used at 10 µM. For the export screen and counter screen, control compounds were used at the following concentrations: dimethyl sulfoxide (DMSO, Thermo Fisher) 0.1% (v/v), Brefeldin A (Sigma Aldrich) 10 µM, Torin 2 (Cayman Chemical) 10 µM, Artemisinin (Sigma) 10 µM, Chloroquine (Sigma) 10 µM. For all other experiments, control compound concentrations were as follows: dimethyl sulfoxide (DMSO, Thermo Fisher) 0.1% (v/v), Brefeldin A (Sigma Aldrich) 5 µg/mL, Torin 2 10 nM. For resistance selection, immunofluorescence assays, microfiltration assays and cytoadhesion assays, further stocks of MMV compounds were sourced from MolPort or MMV and used at 10x their growth EC_50_ as follows: MMV396797 (MolPort-002-021-968): 4.8 µM, MMV665953 (MMV): 20.6 µM.

### Nanoluciferase export assay

Synchronised and drug treated parasites at 1% parasitaemia and 1% haematocrit in complete culture medium were prepared and 5 μL of each culture was added to 8 replicate wells of a 96-well plate. To adjust for the effects of each lysis buffer on nanoluciferase activity, purified recombinant His-nanoluciferase made in house was added to 3D7 parasites not expressing nanoluciferase to 1 ng/mL before 5 uL was added to 8 replicate wells. For differential permeabilization four lysis buffers were prepared as detailed below and 90 μL of each buffer was added to 2 wells of each sample. Tris-phosphate buffer (10 mM Tris-phosphate (Astral Scientific), 132 mM NaCl (Sigma Aldrich), 5 mM EDTA (Astral Scientific), 5 mM DTT (Astral Scientific), pH 7.4) was used as a control for background lysis. EQT buffer (10 mM Tris-phosphate, pH 7.4, 132 mM NaCl, 5 mM EDTA, 5 mM DTT, 4.89 μg/mL purified Equinatoxin made in house was used to selectively lyse the RBC membrane. Saponin buffer (10 mM Tris-phosphate, 132 mM NaCl, 5 mM EDTA, 5 mM DTT, 4.89 μg/mL purified Equinatoxin, 0.03% saponin (Kodak), pH 7.4) selectively lysed the RBC membrane and PVM with Equinatoxin and saponin respectively. Hypotonic buffer (10 mM Tris-phosphate, 5 mM EDTA, 5 mM DTT, 0.2% Igepal CA-630 (Sigma Aldrich), pH 7.4) was used for lysis of all compartments. Finally, 5 μL Tris-phosphate buffer containing 1:50 Nanoglo substrate (Promega) was added to each well. Nanoluciferase activity was measured using a CLARIOStar multi-well plate reader, where samples were shaken at 700 rpm for 30 seconds before luminescence was measured for 1 second per well at 3000 gain. Calculations of relative signal in each compartment and the associated standard deviations were calculated as previously described ^65^.

### Counter screen

Trophozoite stage Hyp1-Nluc parasites at 1% parasitaemia were pelleted by centrifugation at 500g and lysed in 450x pellet volume hypotonic buffer (10 mM Tris-phosphate, 5 mM EDTA, 5 mM DTT, 1x complete protease inhibitor cocktail (Roche) pH 7.4) for 5 minutes at room temperature. Separately, control uninfected RBCs were lysed in 900x pellet volume hypotonic buffer. Samples were centrifuged at 500g for 5 minutes and supernatants were collected. Lysates were combined 1:1 with either hypotonic buffer, or hypotonic buffer with 0.8% Igepal CA-630. Hypotonic lysates (+/- Igepal CA-630) were plated 1:1 with each MMV compound in hypotonic buffer in duplicate to a final concentration of 10 μM for each compound and 0.2% Igepal CA-630 in hypotonic buffer. After a 30-minute incubation at room temperature, 95 uL of each plated sample was transferred to a white luminescence plate and 5 μL Tris-phosphate buffer containing 1:50 Nanoglo substrate was added to each well. Total nanoluciferase bioluminescence was measured using a CLARIOStar multi-well plate reader as described above.

### Western Blotting

Magnet purified PV1-HA-glmS trophozoite-infected or uninfected RBCs were collected by centrifugation at 500g and differentially permeabilized for 5 minutes at room temperature in each of the four lysis buffers detailed previously for the nanoluciferase export assay, plus 1x complete protease inhibitor cocktail (Roche). Non-permeabilised RBCs were pelleted at 500g/4°C/5 minutes and the supernatant fraction was collected and centrifuged again at 3200g/4°C/5 minutes then 16000g/4°C/20 minutes to remove any insoluble material. Supernatant was mixed with NuPAGE sample buffer (Life technologies) to 1x plus 200 mM DTT. Samples were denatured at 70°C for 10 minutes. Proteins were separated by electrophoresis using 4-12% acrylamide bis-tris gels (Invitrogen) in 1x NuPAGE MES SDS running buffer (Invitrogen) for 40 minutes at 200 V. Precision Plus Protein Standard (BioRad) was run as molecular weight marker. After electrophoresis proteins were transferred to a nitrocellulose membrane (BioRad) using an iBlot transfer system (Invitrogen) for 7 minutes. Membranes were then blocked with 1% casein (Sigma Aldrich) in PBS for 1 hour at room temperature before probing with anti-*Hs* haemoglobin (rabbit IgG, 5 µg/mL, Sigma Aldrich), anti-HA (mouse monoclonal Clone HA-7, 10 µg/mL, Sigma Aldrich) or anti-*Pf* ERC (rabbit serum, 1/500, a kind gift from Leann Tilley) primary antibodies in 1% casein/PBS. Blots were washed 3x with PBS and probed with Alexa Fluor 680 Goat Anti Mouse and Goat anti-Rabbit IgG (H+L) Alexafluor Plus 800 secondary antibodies (1/2000, Invitrogen) in 1% casein/PBS and visualized using a LiCor Odyssey infrared imager.

### Immunofluorescence assay

Parasites were synchronised to a 2-4 hour window using percoll density purification and sorbitol lysis and drug treatment was performed for 5 hours at 20-24 (anti-nanoluciferase, anti-HA) or 13-15 (anti-GBP130) hours post invasion. After drug treatment, 15 µL of 4% haematocrit (HCT) culture was allowed to settle onto coverslips coated in 0.1% Poly-L-Lysine (Sigma Aldrich) for 15 minutes and fixed in 4% paraformaldehyde (PFA, Sigma Aldrich) and 0.0075% glutaraldehyde (ProSciTech) in PBS for 20 minutes. After one wash in PBS, cells were permeabilised in 0.1% Triton X-100 (Sigma Aldrich) in PBS for 10 minutes. Cells were washed twice in PBS, 0.02% Triton X-100 and blocked for 1 hour in a 3% Bovine Serum Albumin (BSA, Sigma Aldrich), 0.02% Triton X-100 blocking solution. All antibody incubations were performed in this blocking solution. Cells were probed with primary antibodies overnight at 4°C at the following concentrations: rabbit anti-EXP2 (1:1000, ^43^), mouse anti-GBP130 (1:500), mouse anti-EXP2 (1:5000, ^37^), rabbit anti-Nluc (1:400, ^65^) and mouse anti-HA (1:500, Sigma-Aldrich). After primary antibody was removed, cells were washed 3 times in PBS, 0.02% Triton X-100 and incubated with AlexaFluor 488/594 goat anti mouse/rabbit secondary antibodies (1:2000) with 5% goat serum for 1 hour. Cells were washed 3 times in PBS, 0.02% Triton X-100 and mounted onto slides with Vectashield anti-fade mounting media containing 4’,6-diamindino-2-pheylindole (DAPI, Vector Laboratories). Slides were imaged on a Zeiss Cell Observer widefield fluorescence microscope and analysed using ImageJ.

### Image Analysis

The RBC membrane from the DIC image and PVM from the EXP2 image for each cell were traced using ImageJ and the total fluorescence intensity of exported Hyp1-NLuc or GBP130 within each area was measured for 50 cells in each drug treatment group. For each cell the signal within the PV was subtracted from signal in the whole iRBC to determine exported signal. Mean fluorescence signal was calculated for each compartment and mean exported signal was divided by mean retained signal to obtain a ratio of protein export. These ratios were normalised to the DMSO control group and represented as a percentage.

### Filtration Assay

The relative deformability of infected RBCs was measured using spleen-mimicking microfiltration as previously described ^72^. Synchronous parasite cultures were collected at 14-16 hpi and adjusted to 2-10 % parasitemia, 2% haematocrit in RPMI. Cells were drug treated for 5 hours to 19-21 hpi and passed through a microbead matrix (50% 5-15 µm diameter, 50% 15-25 µm diameter, Industrie des Poudres Sphériques) by adding 800 μL of infected RBC suspension to the column followed by 8 mL RPMI at a flow rate of 0.8 mL/minute. Flow-through samples were collected and centrifuged at 500g for 4 minutes washed in RPMI and centrifuged again at 500g for 4 minutes. Pelleted cells were smeared on a glass slide and parasitaemia was analysed by Giemsa staining. Microfiltration experiments were performed in sets of 3 technical repeats for each biological replicate. Percentage flow-through was calculated by dividing the parasitaemia in the eluted fractions by that of the input fraction and multiplying the result by 100.

### Binding under flow

Ibidi μ-Slide 0.2 channel slides were incubated with 100 μL chondroitin sulfate A (100 μg/mL; Sigma Aldrich) in 1X PBS overnight at 37°C. Channels were blocked with 1% (w/v) BSA– PBS for 1 hour at room temperature and flushed with warm bicarbonate-free RPMI. CS2 parasites panned against Chondroitin Sulfate A ^94^ were synchronised to 15-20 hpi, drug treated for 5 hours to 20-25 hpi and diluted to 3% parasitemia and 1% hematocrit in bicarbonate-free RPMI 1640. Cells were passaged through the channel at 100 μL/min for 10 min at 37°C. Unbound cells were washed from the channel at 100 μL/min for 10 min at 37°C. Bound cells were imaged by widefield microscopy using a Zeiss Cell Observer widefield fluorescence microscope and counted over 10 fields of view (281.28 x 178.14 μm each) using ImageJ.

### Resistance selection

Clonal 3D7 parasites were grown to 5% parasitaemia at 4% haematocrit and 5 mL was added to each well of a 6-well plate (1x 10^8^ parasites per well). MMV396797 was added to 5 wells at 10x the growth EC_50_ (4.8 μM), DMSO was added to the final well at 0.1%. Cultures were maintained at 37°C with new media and MMV396797 or DMSO daily until parasites under MMV396797 selection appeared dead upon inspection by Giemsa smear (3-4 days). Parasites were maintained under regular culture conditions and allowed to replicate and recover. The drug selection process was repeated for a total of 4 cycles.

### Parasite growth assay

Parasite growth was monitored by measuring lactate dehydrogenase (LDH) activity after a 72-hour drug treatment as previously described ^98^. Briefly, synchronous ring-stage 3D7 parasites were diluted to 1% parasitaemia at 1% haematocrit and treated with concentrations of MMV396797 on a 2-step dilution scale from 10 μM to 39 nM for 72 hours at 37°C. Control wells were treated with DMSO at 0.1% (v/v). Samples were then frozen and stored at −80°C before use. After thawing for 4 hours at room temperature, 30 μL of culture from each well was added to 75 μL Malstat mixture comprising a 10:1:1 ratio of Malstat reagent (0.1 M Tris 220 mM lactic acid (Sigma Aldrich), 2% Triton X-100 (v/v), 1.5 mM acetylpyridine adenine dinucleotide (Sigma Aldrich), pH 7.5), 2 mg/mL nitroblue tetrazolium (Sigma Aldrich) and 0.1 mg/mL phenazine ethosulfate (Sigma Aldrich). After mixing samples were incubated in the dark for 45 minutes or until development of colour. Absorbance at 650 nm was measured using a Multiscan Go Microplate Spectrophotometer (Thermo Fisher).

## Acknowledgements

The authors would like to thank the Medicines for Malaria Venture (MMV) for providing access to the MMV Malaria Box. We thank the Australian Red Cross Blood Bank for providing blood and the WEHI Antibody Facility for the EXP2, Nluc and GBP130 antibodies. We thank Matthew Dixon and Leann Tilley for kindly providing CS2 parasites, PV1-HA parasites, microbeads and *Pf* ERC antibodies. We thank Ben Dickerman for providing technical assistance and Thorey Jonsdottir for the LyMP-HA parasites.

---

**Supplementary Table 1. Calculation of Z-scores. A)** The formula for calculating the Z-score between Brefeldin A and DMSO treatment group from the export screen described in Figure 2A. The Z-score is a function of the mean and standard deviation of each exported fraction. Z scores are ranked for separation between means from <0 to 1. **B)** Mean export, standard deviation of mean export, number of total technical repeats and Z-score between DMSO and Brefeldin A treatment groups.

**Supplementary Table 2. Malaria Box export screen.** Full table of export values calculated by screening the entire Malaria Box (as described in Figure 2A-C). Plate coordinates and the MMV number for each compound tested are listed next to the mean (and SEM) Nanoluciferase signal in the exported, secreted, retained and total fractions, presented as a percentage. The total number of technical replicates (n, obtained over multiple biological repeats) is shown, along with a p-value to show which compounds significantly inhibited export compared to the DMSO control (one-way ANOVA, ns p ≥ 0.05, * p < 0.05).

**Supplementary Table 3. Malaria Box counter screen.** A full list of total nanoluciferase signal values calculated by counter screening the Malaria Box (as described in Figure 3A). Plate coordinates and the MMV number for each compound tested are listed with the mean, SEM and SD nanoluciferase signal with or without Igepal CA-630. Values were collected in duplicate from 3 individual repeats (n = 6 total technical replicates). The fold change in signal between MMV and DMSO treatment groups with Igepal CA-630 is listed, where compounds that inhibited nanoluciferase significantly (unpaired t-test, p < 0.05), by >25% or by >50% are listed in separate sheets. The hit compounds from Figure 2B,C and the combined DMSO treatment repeats from each plate are listed in separate sheets.

**Supplementary Figure 1. The Malaria Box contains inhibitors of nanoluciferase. A)** Counter screen of the Malaria Box showing compounds that inhibited nanoluciferase activity by 50% or more. Parasite lysate from Hyp1-Nluc parasites was treated with DMSO (0.1%) or MMV Malaria Box compounds (10 μM) and total nanoluciferase signal was measured. Compounds that inhibited nanoluciferase signal by >50% are indicated in purple. **B)** Nanoluciferase inhibition with and without Igepal CA-630. A list of compounds that inhibited nanoluciferase by >50% from A) showing total nanoluciferase signal in the presence or absence of Ipegal CA-630. Values are shown as mean ± SEM. The statistical difference of the mean for each treatment with or without Igepal CA-630 was calculated using an unpaired t-test (* p < 0.05, ** p ≤ 0.01, *** p ≤ 0.001, **** p ≤ 0.0001). 50% signal is indicated with a dotted line.

**Supplementary Figure 2. Wide field Immunofluorescence microscopy shows retention of LyMP within the bounds of the PV.** Widefield fluorescence microscopy images of LyMP-HA parasites treated with DMSO (0.1 %), Brefeldin A (5 μg/mL), Torin 2 (10 nM), MMV396797 (4.8 μM) or MMV665953 (20.6 μM) for 5 hours from 20-24 hpi. Labelling was performed with rabbit anti-EXP2 ^43^ and mouse anti-HA (Sigma Aldrich) primary antibodies and goat anti-rabbit Alexafluor 594 and goat anti-mouse 488 secondary antibodies. Scale bar = 4 μm.

**Supplementary Figure 3. MMV396797 resistance selection.** Growth of resistance selected parasites in the presence of MMV396797. 3D7 parasites were grown for 72-hours in the presence of MMV396797 at concentrations ranging from 10 μM to 39 nM in a 2-step dilution. Each well represents a 3D7 culture after 4 rounds of drug selection with MMV396797. MMV396797 growth EC_50_ values are indicated in parentheses. Data was collected from 3 technical replicates. Data presented as mean mean ± SD.

**Supplementary Figure 4. MMV665953 strongly inhibits parasite growth.** Light microscopy of Giemsa-stained MMV665953-treated parasites. MSP1-Nluc parasites were treated with DMSO (0.1 %) or MMV665953 (20.6 μM) for 5 hours until 19-21 hpi and Giemsa stained for assessment by bright-field microscopy.

## Notes

### Competing Interest Statement

The authors have declared no competing interest.

