## Supplementary figures and images for "The Medicines for Malaria Venture Malaria Box contains inhibitors of protein secretion in *Plasmodium falciparum* blood stage parasites"

### Supplemental Figure 1

A

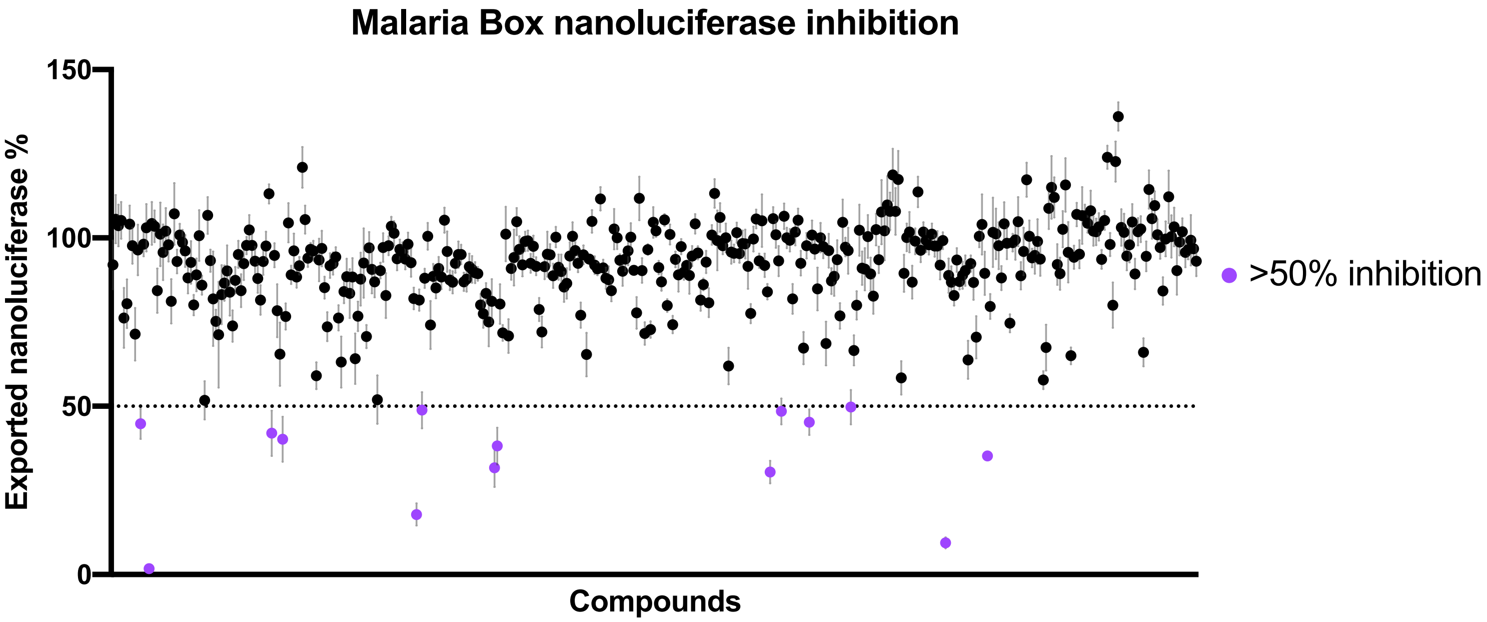


B

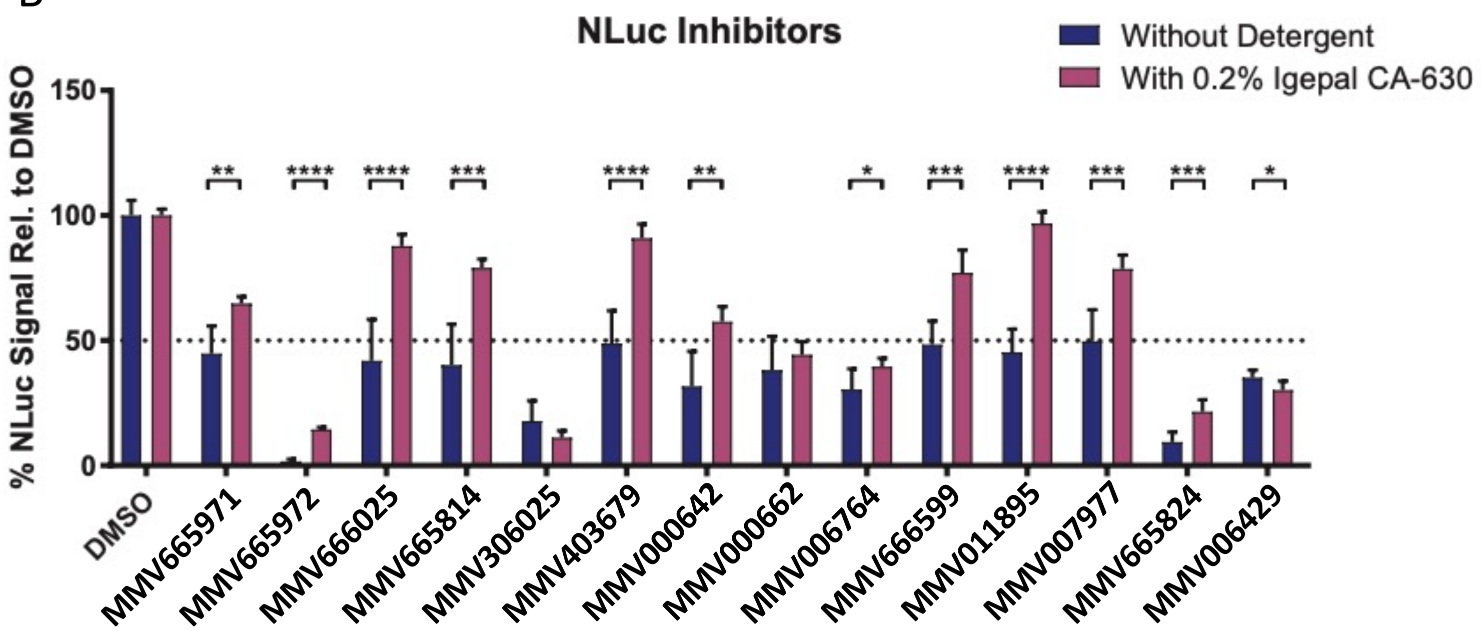


Supplementary Figure 1

### Supplemental Figure 2

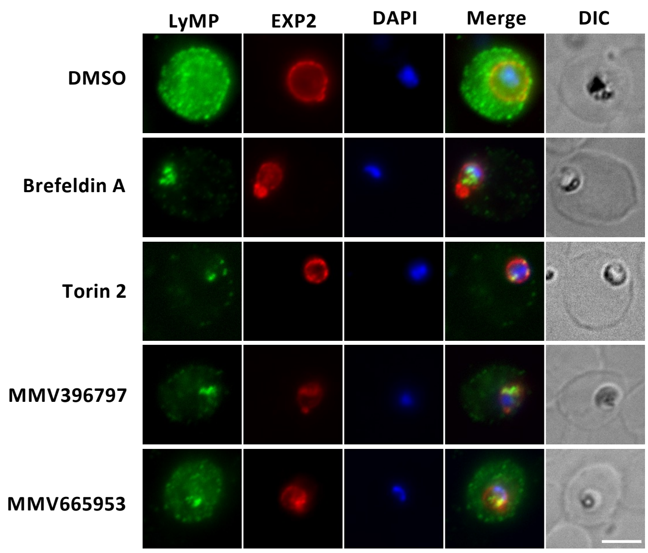


Supplementary Figure 2

### Supplemental Figure 3

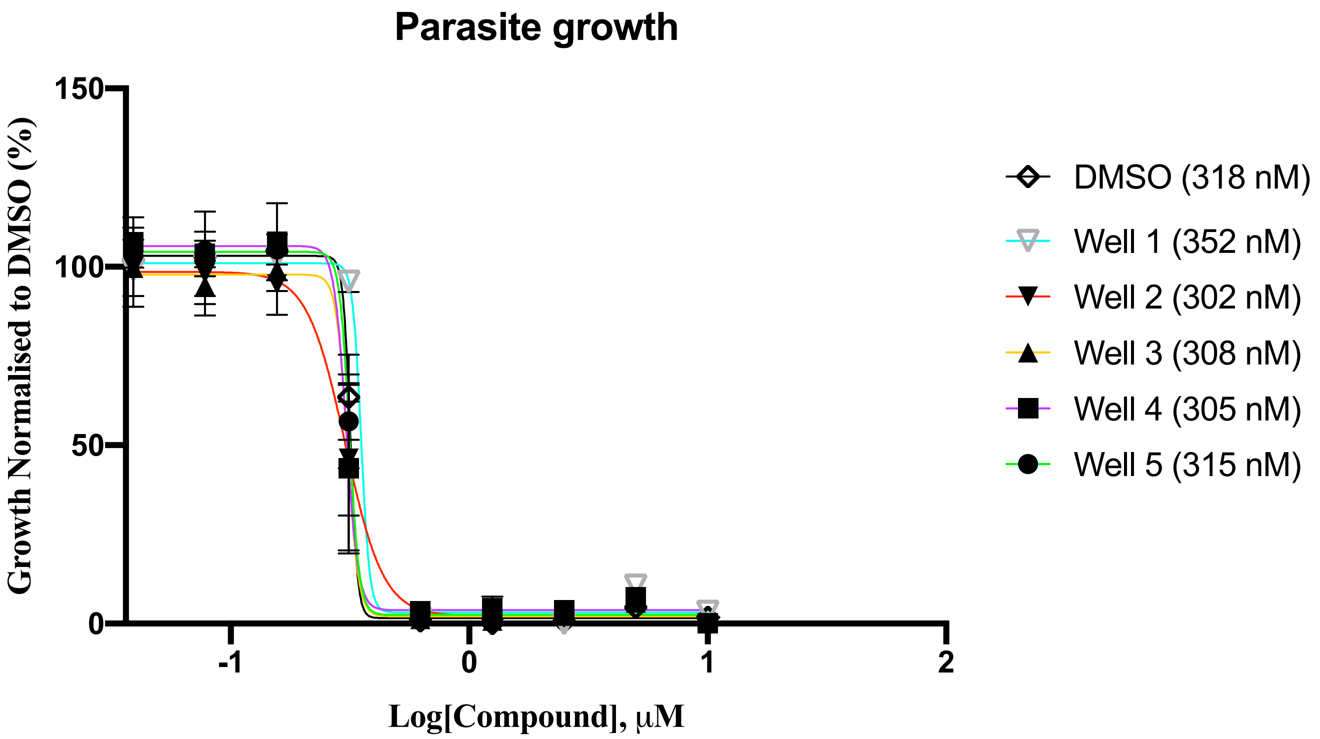


Supplementary Figure 3

### Supplemental Figure 4

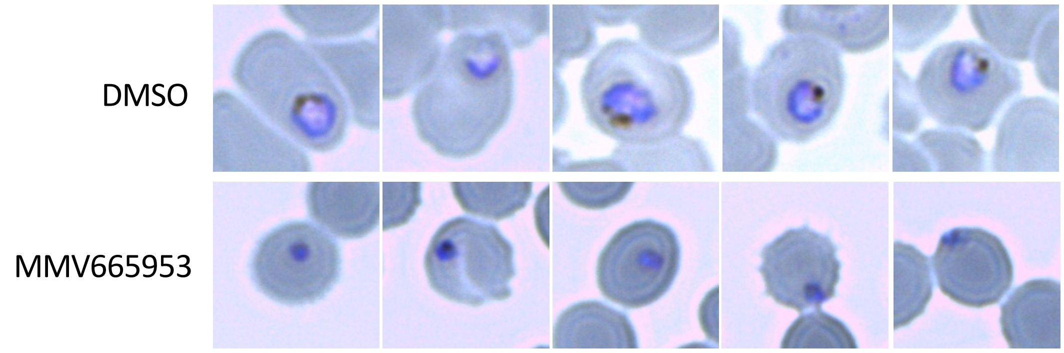


Supplementary Figure 4
