## Supplemental Table 1 for "The Medicines for Malaria Venture Malaria Box contains inhibitors of protein secretion in *Plasmodium falciparum* blood stage parasites"

A

$$Z = 1 - \frac{3S.D._{\text{Brefeldin A}} + 3S.D._{\text{DMSO}}}{|\text{mean}_{\text{Brefeldin A}} - \text{mean}_{\text{DMSO}}|}$$

| **Z-Score** | **Sample separation** |
|---|---|
| **Z = 1** | No variation. SD = 0 |
| **1 > Z > 0.5** | Highly separated sample means |
| **0.5 > Z > 0** | Small separation of sample means |
| **Z < 0** | No separation, variation overlap |

B

| **Treatment** | **Mean export (%)** | **Standard deviation** | **Replicates (n)** | **Z-score** |
|---|---|---|---|---|
| DMSO | 104.9 | 22.2 | 44 | -1.42 |
| Brefeldin A | 59.9 | 14.1 | 20 | |

Supplementary Table 1
